# Kinetic characterization and thermostability of *C. elegans* cytoplasmic and mitochondrial malate dehydrogenases

**DOI:** 10.1101/2021.07.08.451529

**Authors:** Matthew J. Thomas, Emma R. Cassidy, Devin S. Robinson, Katherine M. Walstrom

**Affiliations:** Department of Natural Sciences, State College of Florida, Bradenton, FL 34207, USA; Division of Natural Sciences, New College of Florida, Sarasota, FL 34243, USA

**Author notes:** For correspondence: Katherine M. Walstrom.

**Keywords:** dehydrogenase, enzyme kinetics, protein stability, metabolism, homology modeling, *C*. *elegans*

## Abstract

Malate dehydrogenase (MDH) catalyzes the conversion of NAD^+^ and malate to NADH and oxaloacetate in the last step of the citric acid cycle. Eukaryotes have at least two MDH isozymes, one that is imported into the mitochondria and one that remains in the cytoplasm. We overexpressed and purified *Caenorhabditis elegans* cytoplasmic MDH-1 (F46E10.10) and mitochondrial MDH-2 (F20H11.3) in *E. coli*. Our goal was to compare the kinetic and structural properties of these enzymes because *C. elegans* can survive adverse environmental conditions, such as lack of food and elevated temperatures. In steady-state enzyme kinetics assays, we determined that the *K*_M_ values for oxaloacetate were 54 and 52 μM, and the *K*_M_ values for NADH were 61 and 107 μM, for MDH-1 and MDH-2, respectively. We partially purified endogenous MDH from a mixed population of worms and separated MDH-1 from MDH-2 using anion exchange chromatography. Both endogenous enzymes had a *K*_M_ for oxaloacetate similar to that of the corresponding recombinant enzyme. The reaction velocities of the recombinant enzymes had slightly different temperature-dependencies: MDH-1 and MDH-2 had maximum activity at 40 °C and 35 °C, respectively. In a thermotolerance assay, MDH-1 was much more thermostable than MDH-2. Molecular homology modeling predicted that MDH-1 had more salt-bridges between the subunits than mammalian MDH1 enzymes, and these ionic interactions may contribute to its thermostability. In contrast, the MDH-2 homology model predicted fewer ionic interaction between the subunits compared to mammalian MDH2 enzymes. These results suggest that the increased structural stability of MDH-1 may facilitate its ability to remain active in adverse environmental conditions. In contrast, MDH-2 may use other strategies, such as protein binding partners, to function under similar conditions.

## Introduction

Malate dehydrogenase (MDH^1^) is a central enzyme connecting the citric acid cycle, gluconeogenesis, and the glyoxylate shunt. Eukaryotes contain at least two different isozymes. Mitochondrial MDH2 participates in the citric acid cycle, and cytoplasmic MDH1 has numerous functions because it is an enzyme connecting glycolysis and gluconeogenesis to a variety of reactions occurring in the mitochondria. MDH from many different organisms ranging from archaea to plants to mammals have been studied previously (reviewed in (1–3)). Studies of MDH enzymes from invertebrates have been done, and most of these involved parasitic tapeworms and flukes (4–9). In this study, we characterized *Caenorhabditis elegans* MDH enzymes to determine how the enzymes may be optimized to function in an organism that can withstand intermittent harsh environmental conditions.

Overcrowding and lack of available food can induce *C. elegans* to enter a state of developmental arrest called dauer diapause (reviewed in (10)). Elevated temperatures can also induce dauer formation (11, 12). *C. elegans* can be arrested as dauers for up to 60 days and still recover and return to a normal postdauer lifespan (13). Dauer larvae do not feed, but they remain metabolically active. Specifically, dauers contain increased levels of fat and the protective disaccharide trehalose (1-O-alpha-D-glucopyranosyl-alpha-D-glucopyranoside), and the glyoxylate shunt and fermentation pathways are up-regulated (14–17).

The glyoxylate shunt is in the mitochondria in *C. elegans*. Isocitrate and acetyl CoA molecules enter this pathway to produce succinate and malate via the bifunctional isocitrate lyase/malate synthase enzyme ICL-1 (14, 18–21). Malate can participate in multiple metabolic pathways in the mitochondria or be exported out of the mitochondria. In the latter case, cytoplasmic MDH1 converts malate to oxaloacetate, which enters gluconeogenesis when it is converted to phosphoenolpyruvate (PEP) by PEP carboxykinase. Dauer larvae rely on stored triacylglycerols for energy. Since they are not feeding and consuming sugars, the glyoxylate pathway allows for the degradation of stored lipids to acetyl-CoA and the subsequent production of net oxaloacetate to form glycolytic intermediates and trehalose (14).

The structure, function, and stability of MDH enzymes have been well studied in other other organisms. Numerous crystal structures are available, including structures of thermostable enzymes (22–26). Eukaryotic and bacterial MDH enzymes are usually homodimers, while archaeal MDH enzymes are usually homotetramers (2, 27–31). In this project, we sought to characterize the enzyme kinetic properties of recombinant and endogenous *C. elegans* MDH-1 and MDH-2. We found that the cytoplasmic isozyme, MDH-1, was very thermostable. We then used protein homology modeling to determine possible structural features to explain this stability.

### Experimental Procedures Sequence analysis

The protein sequences from WormBase (wormbase.org) for *C. elegans* MDH-1 (F46E10.10) and MDH-2 (F20H11.3) were aligned with *Sus scrofa* MDH1 (NP_999039.1) and MDH2 (NP_001231082.1), human MDH1 (NP_005908.1) and MDH2 (P40926.3), *Thermus thermophilus* (NCBI:txid274, also called *Thermus flavus*) MDH (P10584.1), and *E. coli* MDH (QIF74064.1) sequences from the National Center for Biotechnology Information (NCBI) using Clustal Omega (32). The MDH-2 sequence was analyzed using MitoProt II v1.101 (33) and MitoFates (34) to determine the cleavage site for the mitochondrial import sequence.

### Plasmid Construction

The *mdh-1* gene was amplified from a mixed population of wild type (Bristol N2) *C. elegans* (NCBI:txid6239) cDNA using primers F46E10_10start, 5’-GGGAATTCCATATGTCCGCCCCACTTCGC - 3’ and F46E10_10end (5’-GGGCAACTAGTGCATCTCCCGTGATGCATGCGATGTTGGCATCATCGCAA GC C- 3’). This PCR product was cut with NdeI and SpeI and ligated into the plasmid pTXB1 (New England Biolabs, NEB) that was also cut with NdeI and SpeI. The sequence of the resulting plasmid was verified by dideoxynucleotide sequencing. The cleaved protein sequence had one extra Ala added to the C-terminus to improve cleavage from the intein tag used for enzyme purification (see below).

The *mdh-2* sequence (with the mitochondrial import sequence removed) was amplified from the cDNA plasmid yk167 (a gift from the Yuji Kohara lab) using the primers MDH_N_Long (5’-GGTTTAGCTAGCAGCCAAGCTCCAAAGGTC) and MDH_C_Long (5’- ATGAACTCGAGGTTTCCCTTAACGAAAGCGACTCCCT).

The PCR product was cut with NheI and XhoI and ligated into plasmid pTXB1 that was previously cut with NheI and XhoI. Then the C-terminal amino acid of MDH-2 was mutated from Ser to Ala to improve the intein cleavage efficiency (see below). This was done using the QuickChange method (Stratagene) and the DNA oligomer 5’- GGAAACCTCGAGGGCTCTG*CG*TGCATCACGGGAG, where the asterisks mark the altered nucleotides that changed the TCC codon to a GCG codon. The mutated plasmid sequence was verified. Recombinant MDH-2 after intein cleavage had an extra MA sequence on the N-terminus and an extra LEGSA sequence on the C-terminus.

### Recombinant Enzyme Expression and purification

Both recombinant enzymes were expressed in freshly transformed Rosetta *E. coli* cells (EMD Biosciences) grown in terrific broth (1.2% tryptone, 2.4% yeast extract, 0.4% glycerol, 0.017 M KH_2_PO_4_, 0.072 M K_2_HPO_4_) and 0.1 mg/mL ampicillin to a density of OD600 = 0.4-0.5. MDH-1 was expressed using 0.05 mM IPTG in cells grown for 20 hr at 25 °C. MDH-2 was expressed using 0.1 mM IPTG in cells grown for 16 hr at 25 °C. The cells were pelleted and stored at -20°C until the enzymes were purified. Both enzymes were expressed as a fusion protein with a self-cleaving intein and a chitin binding domain (CBD) on the C-terminus. The fusion proteins were purified on chitin resin using the IMPACT^TM^ expression system (New England Biolabs) according to the manufacturer’s instructions. The cleavage buffer (20 mM Tris-HCl, pH 8.5, 0.5 M NaCl, 50 mM DTT) for the MDH-1 purification included 0.2% Tween 20 to facilitate elution of the cleaved enzyme, and the cleavage was performed over 96 hours at 25 °C. Then the eluted protein was run over a fresh chitin column to remove remaining MDH-1-CBD fusion protein and additional cleaved CBD (see supporting Figure S1). MDH-2 cleavage was performed in cleavage buffer over 24 hours at 4-6 °C. Protein samples were analyzed using homemade 10-11% Laemmli SDS-PAGE gels stained with Coomassie blue (35, 36). The gels were photographed with a digital camera, and the contrast in the images was improved using “Levels” in Adobe Photoshop 22.4.2. Enzyme fractions with comparable purity were combined and dialyzed into dialysis buffer (30 mM potassium phosphate, pH 7.5, 0.1 M KCl, and 30% glycerol) before storage at -20°C. Protein concentrations were determined using a Coomassie Plus (Bradford) Assay kit (Pierce^TM^). Predicted protein molecular weights, amino acid composition, and theoretical pI values were calculated using the ProtParam tool at Expasy.org. Our gel filtration results indicated that both recombinant MDH-1 and MDH-2 were active as dimers (data not shown).

### Endogenous enzyme purification

*C. elegans* wild type (Bristol N2) hermaphrodites were obtained from the Caenorhabditis Genetics Center and grown on OP50 *E. coli* on egg plates (37) at 20°C. Then they were collected, rinsed extensively in sterile M9 buffer (3 g KH_2_PO_4_, 6 g Na_2_HPO_4_, 5 g NaCl, 1 mM MgSO_4_ per liter) to remove the bacterial food, pelleted, and stored at -80°C. Frozen worms were lysed in 20 mM Tris-HCl, pH 8.0, 150 mM NaCl, 0.1% Tween 20, 1 mM DTT, and 1X Roche Complete ULTRA, EDTA-free protease inhibitors. Lysis was performed on ice using 4 x 10 s, 14 W pulses from a probe sonicator (Fisher Scientific) with 50 s pauses between each pulse. The mixture was spun at 22K rcf at 4 °C for 15 min. To exchange the buffer and remove high molecular weight proteins, the supernatant (clarified lysate) was loaded onto a 7 cm high, 1.5 cm dia. P-100 or P-200 BioGel (BioRad) gel filtration column equilibrated in 50 mM CAPS, pH 9.5 and 10 mM KCl. Fractions containing MDH activity (excluding the first fraction) were loaded onto a 0.3 mL DEAE Sepharose FastFlow (GE Healthcare) column equilibrated with the same buffer. The flow-through (containing MDH-2) was collected, and the column was rinsed with five 0.5 mL additions of the same buffer. Then 50 mM CAPS, pH 9.5 and 50 mM KCl was added to the column to elute MDH-1, and five 0.5 ml fractions were collected. These enzyme samples were used for enzyme kinetics measurements immediately after elution and then dialyzed into the dialysis buffer listed above. Protein concentrations were measured as described above. The rest of the MDH-2 flow-through fractions were mixed with two volumes of 50 mM potassium phosphate, pH 7.5, and loaded onto a 0.4 mL Blue Sepharose column (Sigma-Aldrich) that was equilibrated with the same buffer. This column was washed with 50 mM potassium phosphate, pH 7.5, and 100 mM KCl, and MDH-2 was eluted with 50 mM potassium phosphate, pH 7.5, and 600 mM KCl. A small amount of the eluted sample was diluted 100-fold into enzyme kinetics assays, and the rest was concentrated with a Microcon 50 (Millipore).

The protein samples were analyzed using SDS-PAGE gels and photographed as described above except that a silver stain was used to visualize the proteins (38).

### Enzyme Kinetics

MDH-1 and MDH-2 are classified as EC 1.1.1.37. The reagents were purchased from Sigma-Aldrich or ThermoFisher, and oxaloacetate and NAD^+^ solutions were prepared fresh each day. Enzyme kinetics assays were performed in assay buffer (0.1 M phosphate buffer, pH 7.5, and 0.1 M KCl at 24 °C). This same buffer was used when the temperature was varied. For the kinetics experiments, MDH-1 was stored on ice and MDH-2 was stored in a benchtop cooler that had been cooled to -20 °C. We did not notice a significant decrease in enzyme activity during the experiments. The enzymes were diluted 1:100 into assay buffer at 24 °C. To measure the disappearance of NADH, the solutions were monitored continuously at 340 nm for 30-60 s using an Olis-upgraded Cary 14 or a Cary 300 UV-Vis spectrophotometer, and the slopes of the lines were measured.

Experiments with oxaloacetate varying from 0 – 250 μM included 100 μM NADH. Experiments with NADH varying from 0 - 350 μM included 150 μM oxaloacetate. Some of the measurements at lower [NADH] for MDH-2 (Figure 4C) were performed in 0.1 M phosphate, pH 7.0, and 0.1 M KCl. All enzyme kinetics parameters were determined from non-linear least-squares fits of the data to the Michaelis-Menten equation using RStudio (version 1.4.1106).

The assays at different temperatures were performed in assay buffer containing 150 μM oxaloacetate and 100 μM NADH that had been incubated at the assay temperature for at least 5 min. The cuvette was also pre-incubated at the assay temperature for at least 5 min. For the heat tolerance experiments, the MDH-1 and MDH-2 enzymes were incubated at 55 °C in 50 mM Tris buffer, pH 7.5, and 0.5 mM NADH for the times indicated before they were diluted 100-fold into assay buffer containing 150 μM oxaloacetate and 100 μM NADH at 24 °C.

To calculate specific activities for the endogenous enzymes, the units of enzyme activity were calculated based on the velocities measured in assay buffer containing 150 μM oxaloacetate and 100 μM NADH at 24 °C. For the recombinant enzymes, the specific activities and k_cat_ values were calculated based on the V_max_ values when NADH was varied.

### Protein Molecular Modeling and Structure Analysis

The MDH-1 and MDH-2 sequences were submitted to SWISS-MODEL (39), and the predicted structures were built using ProMod3 (version 3.2.0, (40)) with the crystal structures 1BMD.1.A (25) and 2DFD.1.A (unpublished structure from the Structural Genomics Consortium) as the templates, respectively. The models had GMQE and QMEAN scores (41) of 0.79 and -0.62, respectively, for MDH-1 and 0.78 and -0.04, respectively, for MDH-2. The structure of the MDH-1 model was refined by minimizing the free energy using the YASARA energy minimization server (42). The z-scores for the modeled structures were calculated using the ProSA-web tools (43, 44), the bond and rotamer angles were checked using MolProbity (45, 46), and the models passed the Verify 3D assessment (47–49). The results from these enzyme structure tests are shown in the supporting information Figures S3-S5. UCSF Chimera (version 1.15.0) (50) was used to search for intersubunit hydrogen-bonds (with constraints relaxed by 0.4 Å and 20 degrees) involving charged amino acids. UCSF Chimera was also used to search for intersubunit hydrogen bonds in 5MDH (51) and 1BMD (25), and these results corresponded to the published lists of intersubunit hydrogen bonds in both structures (22, 25). The structures of *S. scrofa* MDH2 (1MLD) and an unpublished human MDH2 structure (4WLU) were obtained from the PDB and analyzed for intersubunit hydrogen bonds, and the *S. scrofa* MDH2 interactions mostly agreed with published results (23), as explained in Table 5. The PyMOL Molecular Graphics System (Version 2.4.2, Schrödinger, LLC) was used to measure distances, visualize the subunit interactions, and produce the structure figures. Amino acid composition was analyzed using the ProtParam tool at ExPASy.org. The protein structure volumes were measured using UCSF Chimera with the default settings. See the supporting information for additional protein modeling experimental details.

## Results

### *C. elegans* MDH-1 and MDH-2 sequence analysis

The cytoplasmic MDH in *C. elegans* (F46E10.10) was identified by Holt and Riddle (52), and we identified the mitochondrial enzyme (F20H11.3) by sequence homology. These enzymes were named MDH-1 and MDH-2, respectively, to conform with the naming convention used for MDH enzymes in other eukaryotes. An alignment of the *C. elegans* enzymes with *Sus scrofa* MDH1 and MDH2, human MDH1 and MDH2, *Thermus thermophilus* MDH, and *E. coli* MDH is shown in Figure 1 with a phylogenetic tree and sequence identity tables. As expected, the mitochondrial enzymes were most similar to each other, as were the cytoplasmic enzymes. The *T. thermophilus* sequence was more similar to the cytoplasmic enzymes, and the *E. coli* sequence was more similar to the mitochondrial enzymes, and both of these results have been discussed previously (1, 2).

**Figure 1.**
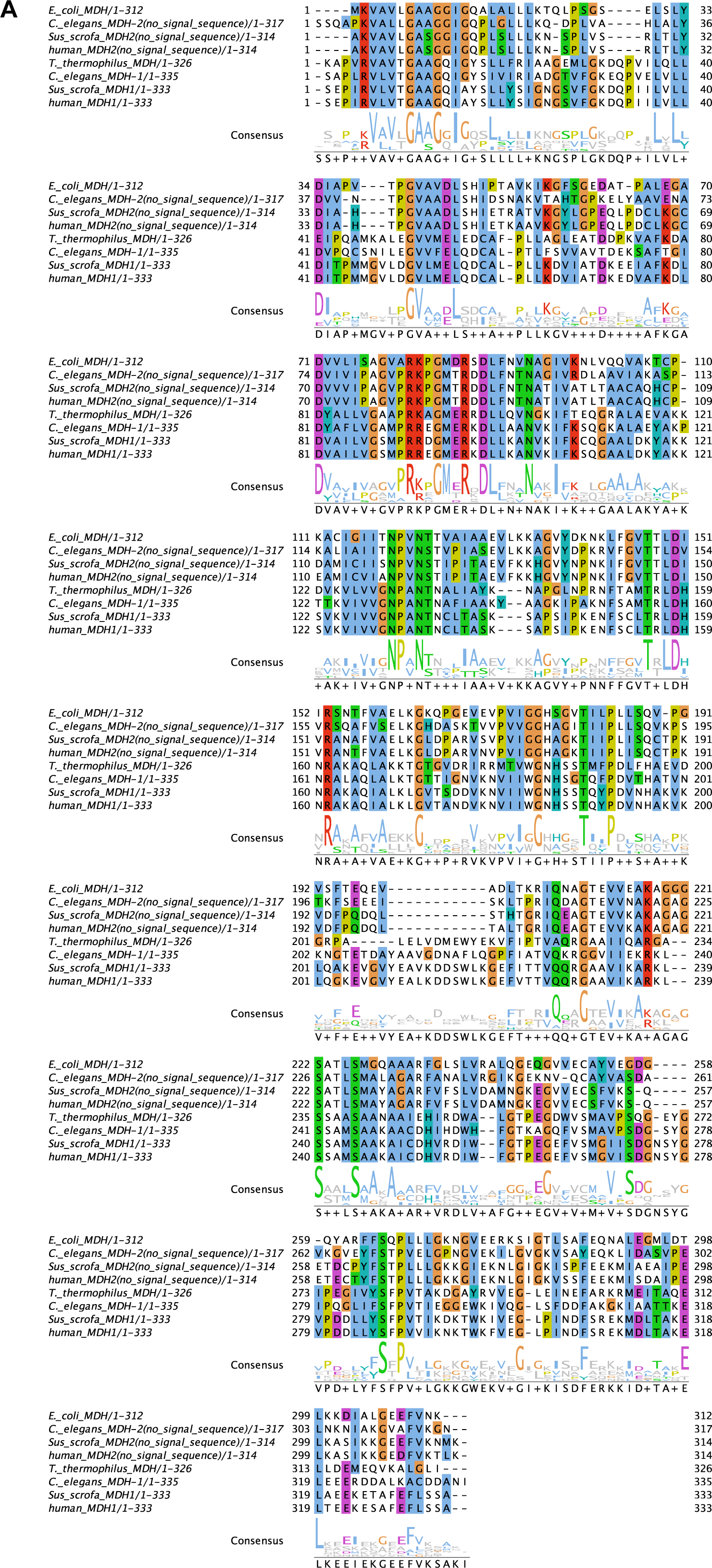
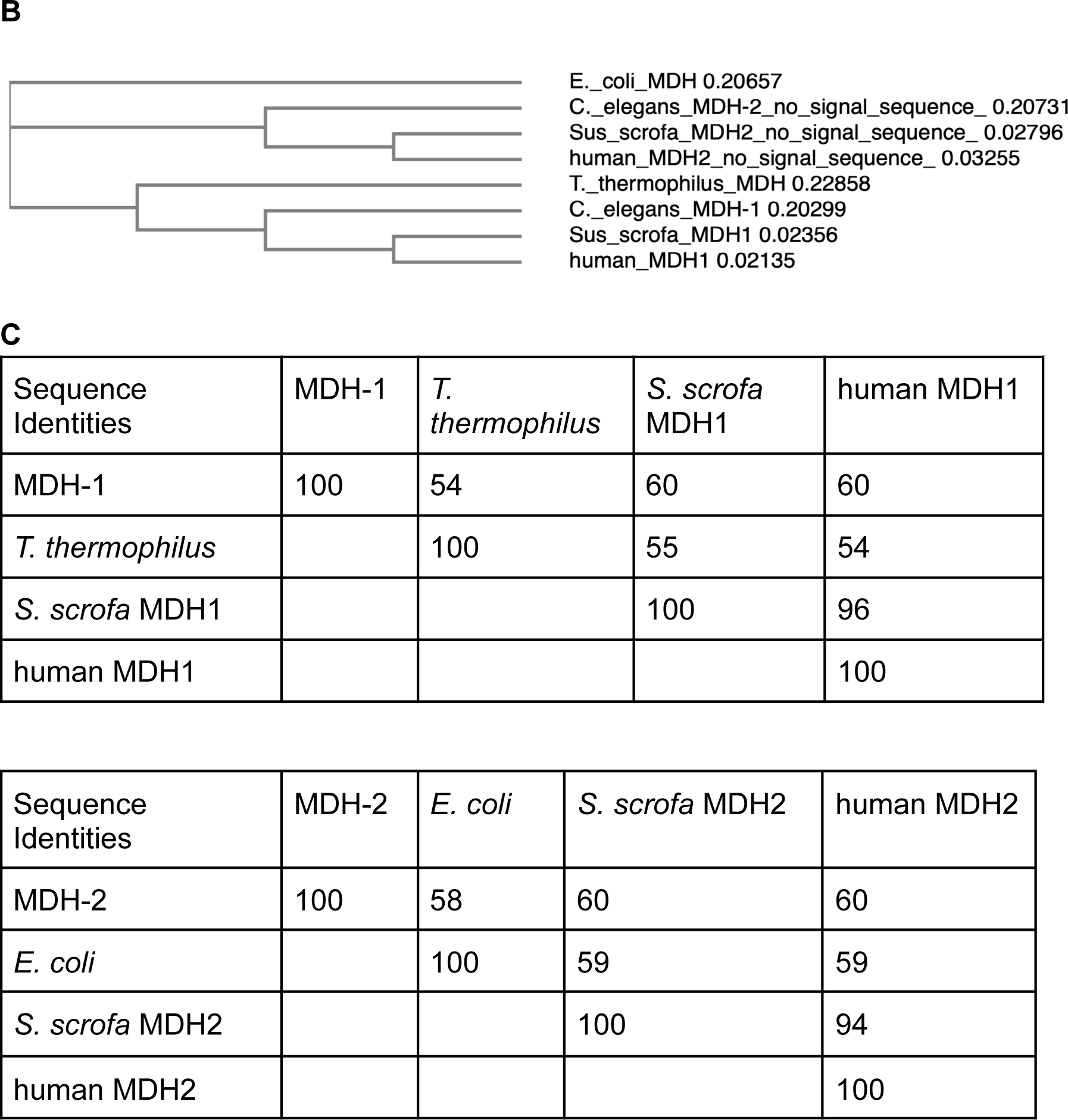
MDH sequence alignments and analysis. **A.** The protein sequences for *C. elegans* MDH-1 (F46E10.10) and MDH-2 (F20H11.3) were aligned with the *S. scrofa* MDH1 (NP_999039.1) and MDH2 (NP_001231082.1), human MDH1 (NP_005908.1) and MDH2 (P40926.3), *T. thermophilus* MDH (P10584.1), and *E. coli* MDH (QIF74064.1) using Clustal Omega (32) and visualized using Jalview (115). Amino acid coloring is based on sequence identity. The MDH1 amino acid numbering starts with the first amino acid after the methionine (which is not shown in the sequences), and the MDH2 numbering starts with the first amino acid after cleavage of the mitochondrial import signal sequence. **B.** The phylogenetic tree for these proteins is a neighbor-joining tree without distance corrections made with Clustal Omega. **C.** The amino acid sequence identities were calculated by Clustal Omega.

The amino acid numbering in Figure 1 corresponds to previous publications. Specifically, the papers describing the structures of *S. scrofa* MDH1 and *T. thermophilus* MDH both used the amino acid numbering of Birktoft, et al. (1989), which started at number 1 for the second amino acid of *T. thermophilus* MDH (because the *S. scrofa* MDH1 lacks the methionine) (22, 25). The *C. elegans* MDH-1 amino acid numbers in this paper follow the *S. scrofa* MDH1 numbering in Figure 1, even though MDH-1 goes out of register by one amino acid between amino acids 142-275 because of an amino acid deletion and addition in two exterior loops (data not shown). The paper describing the *S. scrofa* MDH2 structure started numbering at the first amino acid of the cleaved protein (23), and this convention is used here for all the mitochondrial MDH proteins. Since MDH-2 has 4 extra amino acids on its N-terminus compared to the other MDH2 enzymes, the amino acid numbering for *S. scrofa* was used for MDH-2 and the mammalian MDH2 enzymes.

The substrate-binding and active site residues that have been previously identified are conserved in both *C. elegans* MDH-1 and MDH-2 (22–25, 51, 53). The active site includes (using the MDH1 amino acid numbering in Figure 1) Asn130, His186 (His176 in MDH2), Asp158, and Arg161 as well as Arg91 and Arg97 in the mobile loop that closes over the active site. In MDH1, the adenine ring of NADH binds to Leu40, Ile42, Met45, Thr9, Gly10, Ala12, and Gly13 in a hydrophobic pocket. In MDH1 only, Glu73 hydrogen bonds to N6A of adenine. In *E. coli* MDH, Tyr33, Ile35, Ala77, Ile97, and Leu101 form the hydrophobic adenine binding pocket, and these residues surround the adenine ring in the unpublished human MDH2 structure (4WLU, data not shown). There are hydrogen bonds between the NADH pyrophosphate and Gln14 and Ser240 (neither conserved in MDH2) as well as the backbone of residues of Gln14 (Gly11 in MDH2) and Ile15. Asn130 makes hydrogen bonds to the nicotinamide ribose hydroxyl groups, and Asp41 hydrogen bonds to the hydroxyl groups in the adenine ribose. In MDH1, the residues forming a binding site for the nicotinamide ring include Ile15, Val128, Leu154, Leu157, and Ala245. In MDH2, the backbone oxygen of Ile117 hydrogen bonds with the nicotinamide carboxyamide group.

Numerous studies have shown that mitochondrial MDH2 enzymes interact with other proteins to channel substrates between enzymes (54–58). For example, the interaction between beef heart MDH2 and citrate synthase (CS) involved amino acids surrounding the MDH2 active site opening (58), which are nearly all conserved in *C. elegans* MDH-2, *S. scrofa* MDH2 and human MDH2.

### MDH-1 and MDH-2 Purifications

The recombinant MDH-1 and MDH-2 enzymes were overexpressed in *E. coli* as fusion proteins with a self-cleaving intein and a chitin binding domain (CBD). The MDH-1-CBD and MDH-2-CBD fusion proteins were purified using chitin resin, and the CBD tag was removed from the enzymes using intein cleavage induced by DTT addition. The purification of MDH-2 is shown in Figure 2, and the purification of MDH-1 followed a similar procedure (see the Experimental section and supporting Figure S1). The final recombinant MDH-1 preparation is shown in Figure 3B, lanes 8 and 10, and the final recombinant MDH-2 sample is shown in Figure 1, lanes 7-10, Figure 2A, lane 5, and Figure 2B, lane 6. The predicted molecular weights for the recombinant MDH-1 and MDH-2 enzymes were 35.9 kDa and 33.3 kDa, respectively. The molecular weights based on the size standards in the SDS-PAGE gel in Figure 2 were 33 +/- 3 kDa and 31 +/- 2 kDa (95% confidence intervals) for MDH-1 and MDH-2, respectively.

**Figure 2.**
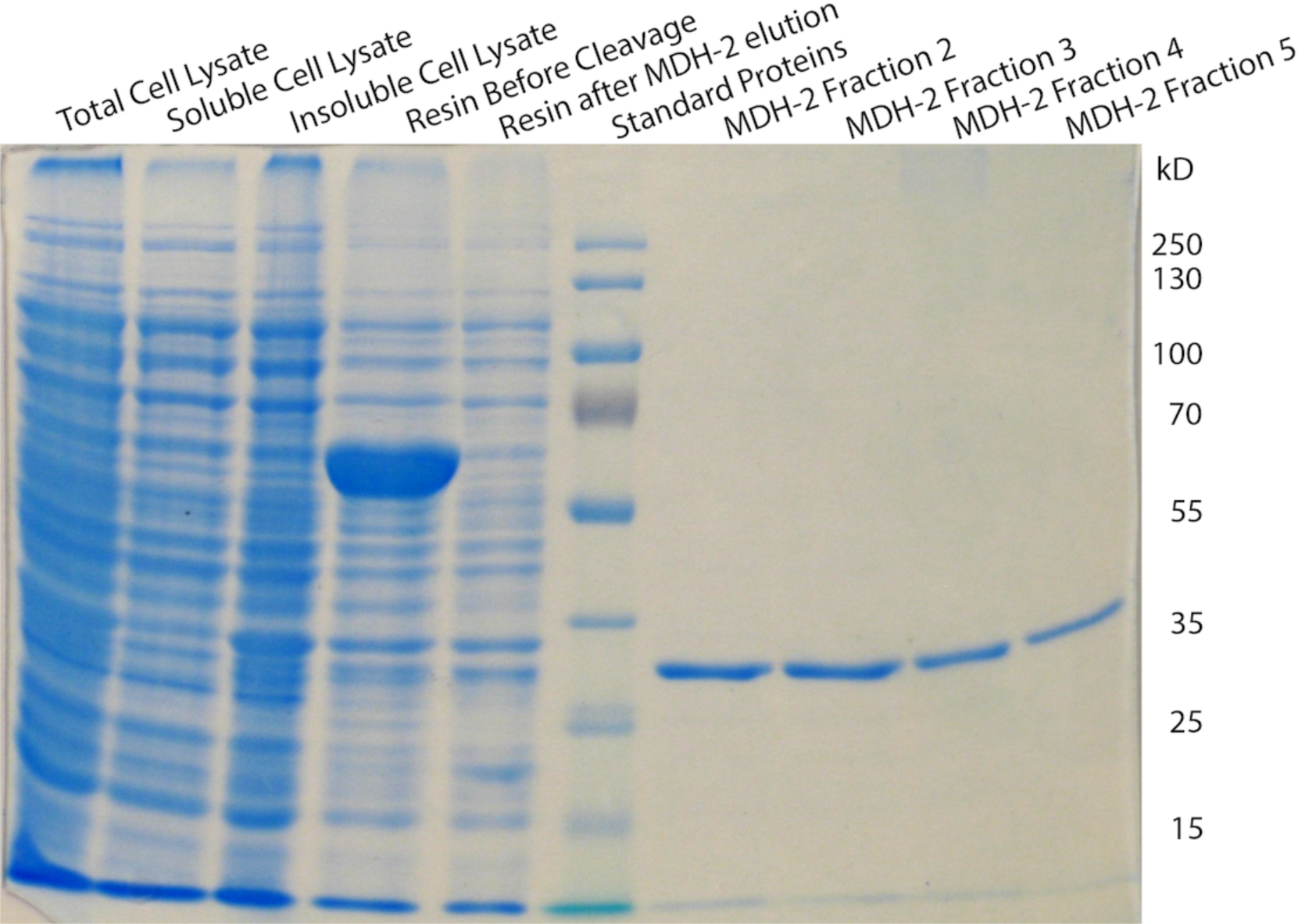
Recombinant MDH-2 purification. Samples from the purification of recombinant MDH-2 are shown in a 10% SDS-PAGE gel. The soluble cell lysate containing the MDH-2-CBD fusion protein (61.2 kDa) was run through a chitin resin column, and the protein bound to the resin (lane 4). Then DTT was added to the resin for 24 hr. at 4 °C to induce cleavage of the intein to release MDH-2 (33.3 kDa). The MDH-2 fractions (lanes 7-10) show that a single polypeptide was released from the resin. The standard protein sizes are shown in kDa on the right side of the gel.

**Figure 3.**
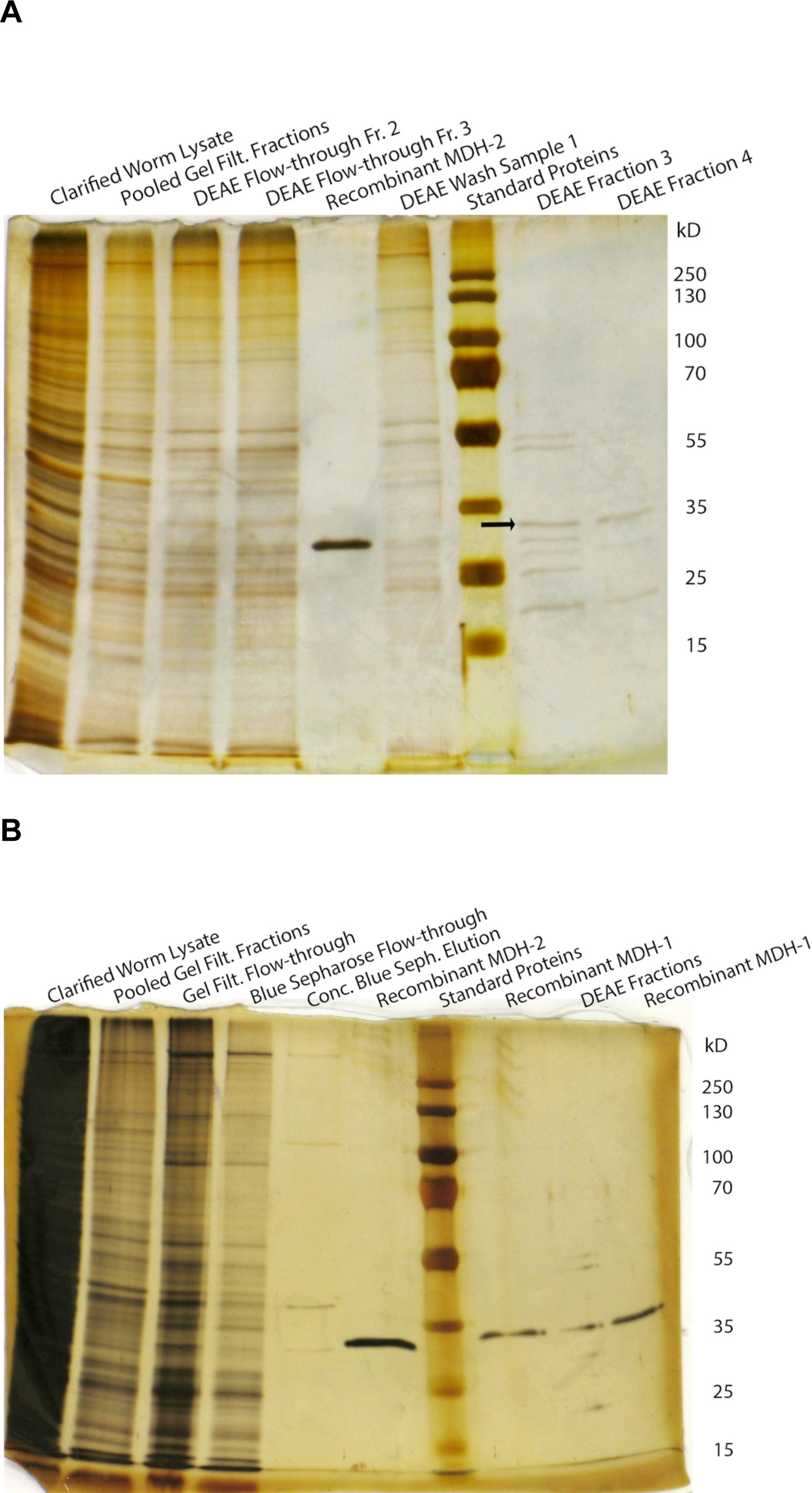
Endogenous MDH-1 and MDH-2 purification. **A.** Samples from a partial purification of endogenous MDH-1 and MDH-2 are shown in a 11% SDS-PAGE gel. The clarified worm lysate was run through a gel filtration column to change the buffer to 50 mM CAPS, pH 9.5, and 10 mM KCl. The gel filtration fractions were loaded onto a DEAE column equilibrated in the same buffer. The flow-through samples from the DEAE column contained MDH-2, and then the KCl concentration was increased to 50 mM in the same buffer to remove MDH-1 from the column. Lanes 8 and 9 show the MDH-1-containing samples with MDH-1 labeled with an arrow. The sizes of the protein standards are shown in kDa. **B.** Samples from a second partial purification of endogenous MDH-1 and MDH-2 are shown in a 10% SDS-PAGE gel. The purification was performed in the same way as shown in 3A. However, the DEAE flow-through samples were loaded onto a Blue Sepharose column in 50 mM potassium phosphate, pH 7.5, to further purify MDH-2. The final MDH-2 sample was eluted with 600 mM KCl in the same buffer and is shown in lane 5 next to recombinant MDH-2 in lane 6. The endogenous MDH-1 sample (fractions 3-5 combined from the DEAE column) is shown in lane 9 between two samples of endogenous MDH-1 in lanes 8 and 10. The sizes of the protein standards are shown in kDa.

The endogenous MDH-1 and MDH-2 enzymes were purified from a *C. elegans* sample that contained worms of various ages. The theoretical pI values for MDH-1 and MDH-2 (after signal sequence cleavage) were 6.6 and 9.1, respectively. The two proteins were separated using DEAE ion exchange, and then MDH-2 was further purified using a Blue Sepharose column (Figure 3B). The purification process is shown in Table 1.

**Table 1.** Purification table for endogenous MDH-1 and MDH-2

| Sample | Total Protein (mg) | Total Units (μmol/min) | Specific Activity | Yield (%) | Purification Level |
| --- | --- | --- | --- | --- | --- |
| Clarified Worm Lysate | 3.9 ± 0.2<sup>1</sup> | 13 ± 1 | 3.3 ± 0.3 | 100 | 1 |
| Gel Filtration Samples (with MDH-1 and MDH-2) | 1.2 ± 0.2 | 6.63 ± 0.2 | 5.4 ± 0.8 | 52 | 1.6 |
| dialyzed DEAE fractions (with MDH-1) | 0.06 ± 0.01 | 1.6 ± 0.1 | 25 ± 5 | 12.6 | 7.6 |
| DEAE Flow-through (with MDH-2) | 0.56 ± 0.02 | 1.11 ± 0.04 | 2.0 ± 0.1 | 8.7 | 0.6<sup>2</sup> |
| Blue Sepharose fractions (with MDH-2) | 0.00062 ± 0.00008 | 0.010 ± 0.001 | 21 ± 3 | 0.1 | 6.4 |

<sup>1</sup>Standard errors are shown.
<sup>2</sup>This value is less than 1.0 because the total number of units in the worm lysate included the activity of both MDH-1 and MDH-2. This DEAE flow-through sample only contained activity from MDH-2.

Samples from the endogenous protein purification were run on SDS-PAGE gels next to the recombinant proteins, and there were bands of the expected size in each endogenous sample. The final preparation of endogenous MDH-1 contained about 6 proteins, and the band at the expected size for MDH-1 was the most intense band (Figure 2B, lane 9). The final preparation of MDH-2 contained about 4 proteins, and the band at the expected size for MDH-2 was less intense than some of the other bands (Figure 2B, lane 5). The predicted molecular weights for endogenous MDH-1 and MDH-2 were 35.8 kDa and 32.7 kDa. The molecular weights based on the size standards in the SDS-PAGE gel in Figure 2B were 33 +/- 3 kDa and 30 +/- 2 kDa (95% confidence intervals) for MDH-1 and MDH-2, respectively.

Based on the units (U, μmol/min) of enzyme activity of the endogenous MDH-1 and MDH-2 samples used for enzyme kinetics (Table 1), the *C. elegans* sample contained slightly more MDH-1 activity (1.6 U) than MDH-2 activity (1.1 U). This is almost the same ratio (1.5) of MDH activity in the cytoplasmic fraction compared to the mitochondrial fraction from adult *C. elegans* measured in a previous study (19). The specific activities of the final endogenous MDH-1 and MDH-2 samples were similar (Table 1), but much more of the MDH-2 was lost during the purification because the SDS-PAGE sample (Figure 2B, lane 5) went through an extra Blue Sepharose column and ultrafiltration step.

### Enzyme Kinetics

Steady-state enzyme kinetics measurements were performed with the purified endogenous and recombinant MDH-1 and MDH-2 enzymes and analyzed using the Michaelis-Menten equation (59, 60). Since the forward MDH reaction is energetically unfavorable at neutral pH, the reverse reaction is usually studied *in vitro* (2, 61). The pH-dependence of MDH-2 activity was tested from pH 4 - 11 (see supporting Figure S2), and the enzyme activity was high between pH 6 - 8.5 in phosphate and Tris-HCl buffers. Figures 4A and 4B show the enzyme kinetics results when oxaloacetate was varied, and Figure 4C shows the results for the recombinant enzymes when NADH was varied. The *K*_M_ and V_max_ values are shown in Table 2. Blank samples containing all of the reaction components except oxaloacetate were tested for every enzyme preparation, and the rate of change of the absorbance at 340 nm was at least 10-fold lower than the rates measured at the lowest oxaloacetate concentrations.

**Figure 4.**
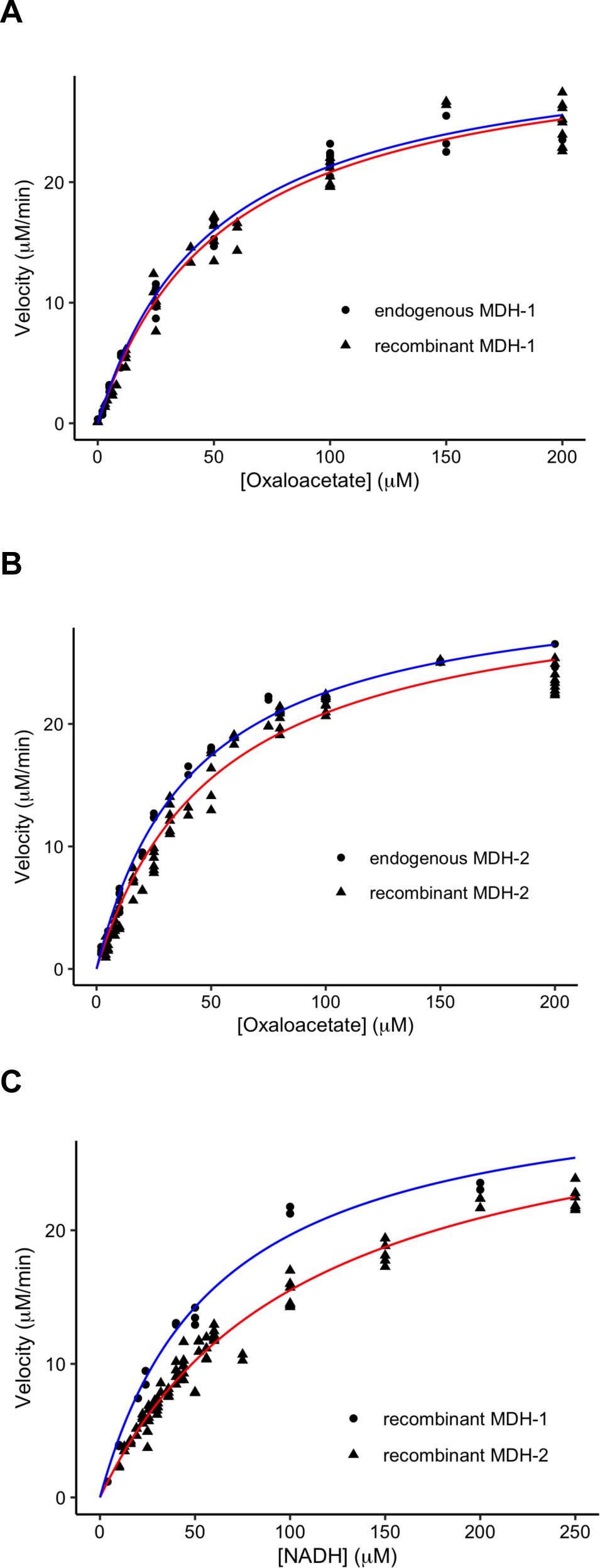
MDH-1 and MDH-2 Enzyme Kinetics. Enzyme velocities of recombinant (triangles) and endogenous (circles) MDH-1 (**A**) and MDH-2 (**B**) are shown as oxaloacetate was varied. The rate of change of absorbance at 340 nm was monitored as NADH was covered to NAD^+^ during the reaction, and this was converted to μM/min. All of the data points are shown from replicate experiments performed at least twice. For these graphs, the data were normalized so that the V_max_ values were the same for each enzyme. The data were fit to the Michaelis-Menten equation, and the best-fit lines are shown in red and blue for recombinant and endogenous enzymes, respectively. The reaction conditions were 0.1 M sodium phosphate buffer, pH 7.5, 0.1 M KCl, 100 μM NADH, and oxaloacetate was varied from 0 to 200 μM at 24 °C. **C.** Enzyme velocities of recombinant MDH-1 (circles, blue line) and MDH-2 (triangles, red line) are shown as NADH was varied. The procedures were the same as described for 4A and 4B, except that 150 μM oxaloacetate was used, and the NADH concentration was varied from 0 to 250 μM.

**Table 2.** Summary of Michaelis-Menten values

| Enzyme | $K_M$, $\mu\text{M}$ | $K_M$ std error | $V_{\text{max}}$, $\mu\text{M}/\text{min}$ | $V_{\text{max}}$ std error |
| --- | --- | --- | --- | --- |
| Values when oxaloacetate was varied |  |  |  |  |
| Endo. MDH-1 | 50 | 4 | 26.5 | 0.8 |
| Recomb. MDH-1 | 54 | 4 | 32 | 1.0 |
| Endo. MDH-2 | 42 | 2 | 22.1 | 0.5 |
| Recomb. MDH-2 | 52 | 4 | 66 | 2 |
| Values when NADH was varied |  |  |  |  |
| Recomb. MDH-1 | 61 | 6 | 40 | 2 |
| Recomb. MDH-2 | 107 | 5 | 113 | 3 |

The *K*_M_ values for recombinant and endogenous MDH-1 were quite similar: 54 ± 4 μM and 50 ± 4 μM, respectively, with the standard errors shown. The *K*_M_ values for recombinant and endogenous MDH-2 varied more: 52 ± 4 μM and 42 ± 2 μM, respectively. This difference may be because the recombinant MDH-2 enzyme had a few extra amino acids on the N- and C-termini (see Experimental Methods). The *K*_M_ values for NADH were 61 ± 6 μM and 107 ± 5 μM for recombinant MDH-1 and MDH-2, respectively. The *K*_M_ values for NADH for the endogenous enzymes were not determined.

We attempted to do enzyme kinetics with malate and NAD^+^ with recombinant MDH-1 and MDH-2 under a variety of reaction conditions. Specifically, we tried higher concentrations of each reagent and higher pH values, because these conditions have worked for some MDH enzymes (5, 6, 62). However, we were not able to observe any enzyme activity with malate and NAD^+^ under any of the conditions we tried (data not shown).

The protein concentrations and the V_max_ values when NADH was varied were used to calculate the specific activities and k_cat_ values for the recombinant enzyme preparations (Table 3). As shown in the recombinant MDH-1 and MDH-2 purification gels (Figure 1 and supporting Figure S1), the MDH-1 enzyme expressed to a higher level in *E. coli*. The final specific activity of recombinant MDH-2 was slightly higher than that for MDH-1 (Table 3). This may be because the MDH-1 preparation contained a low level of contaminating MDH-1-CBD fusion protein and cleaved CBD (see supporting Figure S1).

**Table 3.** Specific Activities and k_cat_ values of recombinant MDH-1 and MDH-2

| Sample | Total Protein (mg) | Total Units | Specific Activity, U/mg | $k_{cat}$, $s^{-1}$ |
| --- | --- | --- | --- | --- |
| MDH-1<sup>a</sup> | $2.01 \pm 0.07^c$ | $1190 \pm 57$ | $590 \pm 35$ | $350 \pm 21$ |
| MDH-2<sup>b</sup> | $0.27 \pm 0.01$ | $227 \pm 6$ | $828 \pm 41$ | $460 \pm 23$ |

<sup>a</sup>Measured using the combined fractions 1 and 2 for MDH-1 and calculated using the $V_{max}$ when NADH was varied.
<sup>b</sup>Measured using the combined fractions 1 and 2 for MDH-2 and calculated using the $V_{max}$ when NADH was varied.
<sup>c</sup>Standard errors are shown.

The protein concentrations and the reaction rates measured at 150 μM oxaloacetate and 100 μM NADH were used to calculate the specific activities of the endogenous enzymes (Table 1). The enzyme kinetics for endogenous MDH-1 was measured using the DEAE fractions, shown in lanes 8 and 9 in Figure 3A and lane 9 of Figure 3B. The enzyme kinetics for endogenous MDH-2 was measured using the DEAE flow-through fractions, such as those shown in lanes 3 and 4 in Figure 3A. The MDH-2 sample after the Blue Sepharose column that is shown in Figure 3B, lane 5 was used to determine a specific activity of 21 U/mg, compared to 25 U/mg for MDH-1 (Table 1). These results are fairly consistent with the SDS-PAGE gels (Figure 3B), because the MDH-1 band is more intense compared to the contaminating protein bands, while the MDH-2 band is faint compared to the contaminants.

### MDH-1 and MDH-2 Thermostability

During our experiments, we noticed that MDH-1 was very stable, even when left at room temperature for a couple days. To test whether recombinant MDH-1 and MDH-2 have different temperature optima, we measured the temperature-dependence of the activity of each enzyme (Figure 5). The optimum temperature of MDH-1 was about 40 °C, while the optimum for MDH-2 was about 35 °C. To determine the thermostability of the enzymes without any interference from the temperature-sensitivity of the substrates, we incubated the recombinant enzymes at 55 °C for different lengths of time and then measured the residual enzyme activity at 24 °C (Figure 6). This experiment showed a clear difference in the thermostabilities of MDH-1 and MDH-2. MDH-1 maintained its activity for about 10 min at 55 °C, while MDH-2 lost its activity in about 1 min. The thermostability of MDH-1 was interesting because *C. elegans* have an optimum growing temperature in the laboratory of 20 °C, and temperatures above 28 °C can cause detrimental effects such as sterility (63, 64).

**Figure 5.**
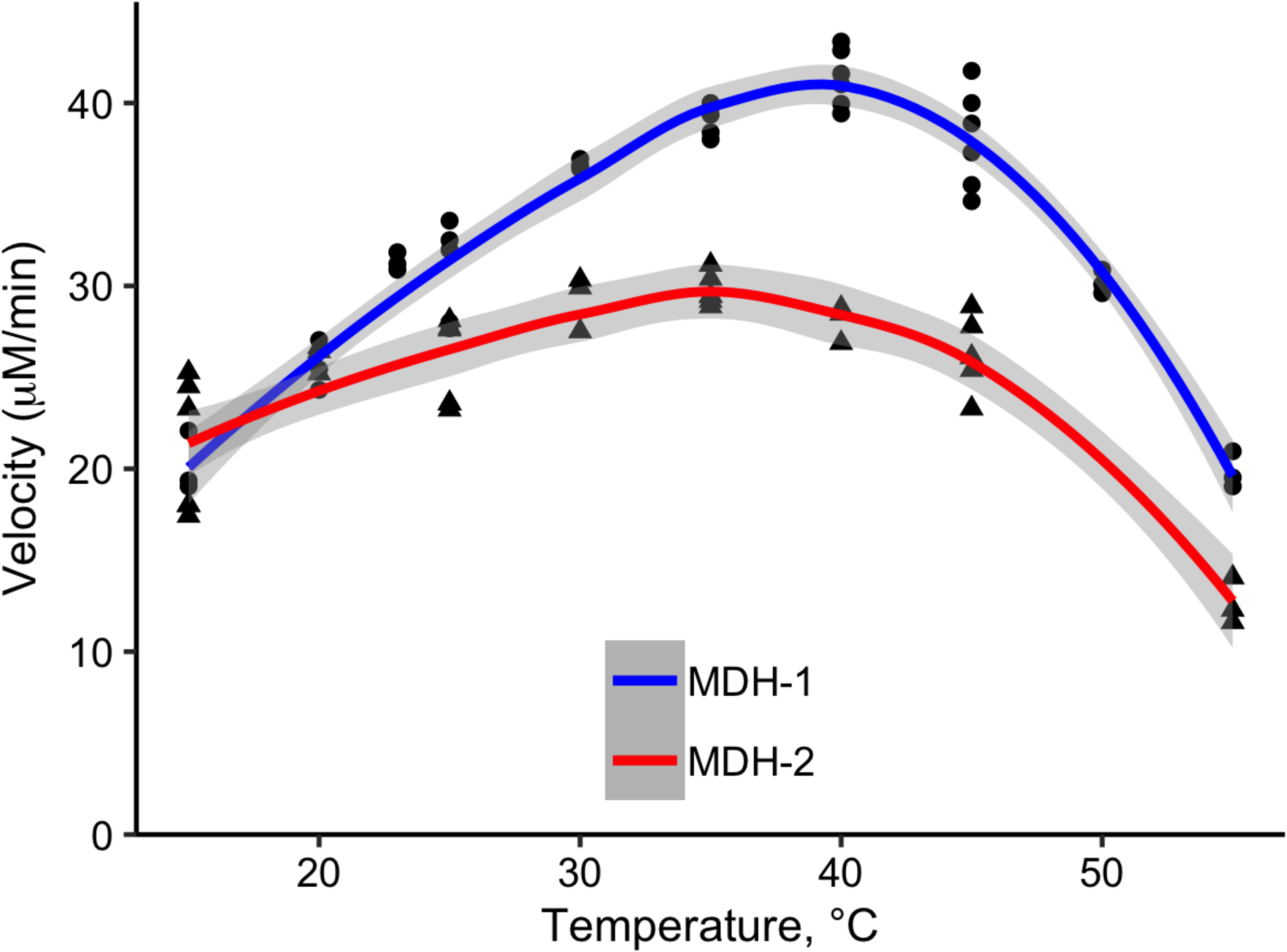
Temperature-dependence of MDH-1 and MDH-2 enzyme activity. Enzyme kinetics experiments were performed with MDH-1 (circles) and MDH-2 (triangles) at temperatures from 15 to 55 °C. The reaction conditions were 0.1 M sodium phosphate buffer, pH 7.5, 0.1 M KCl, 100 μM NADH, and 150 μM oxaloacetate. The data were smoothed using the Loess method, and the best-fit lines (blue for MDH-1 and red for MDH-2) and 95% confidence intervals (gray) are shown. All of the data points are shown from replicate experiments performed at least twice. The velocity values were normalized using the average value at 20 °C.

**Figure 6.**
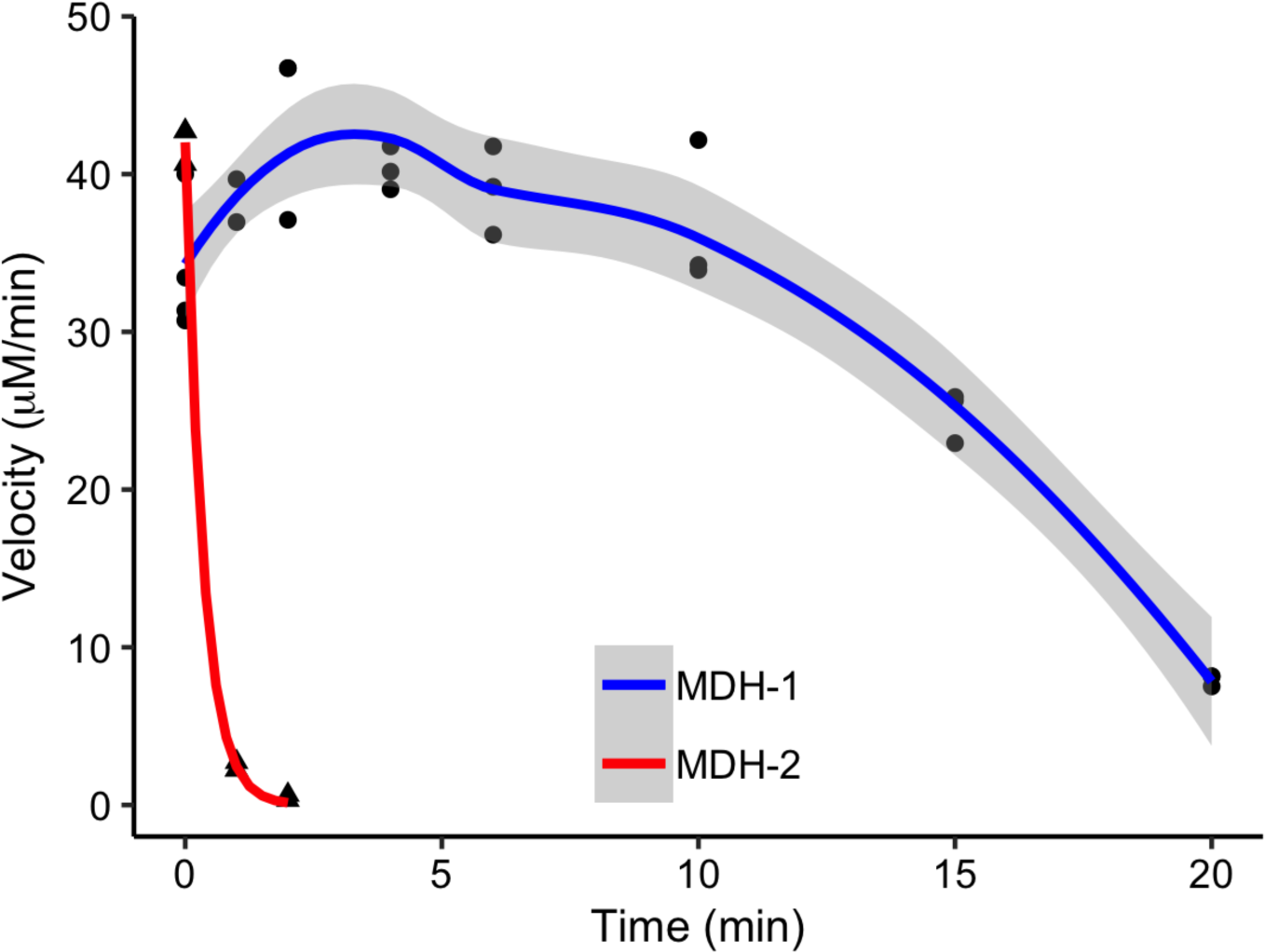
Thermostability of recombinant MDH-1 and MDH-2. Recombinant MDH-1 and MDH-2 in 50 mM Tris buffer, pH 7.5, and 0.5 mM NADH were placed in a water bath at 55-57 °C for times ranging from 0 - 20 min. At each time point, enzyme samples were taken and used immediately for enzyme kinetics measurements. The enzyme kinetics reaction conditions were 0.1 M sodium phosphate buffer, pH 7.5, 0.1 M KCl, 100 μM NADH, 150 μM oxaloacetate, and 24 °C. The data were smoothed using the Loess method (for MDH-1) and fit to an exponential (for MDH-2). The best-fit lines (blue for MDH-1 and red for MDH-2) and the 95% confidence interval for MDH-1 (gray) are shown. The rate constant for inactivation of MDH-2 was 2.85 ± 0.18 min^-1^ (st. error). All of the data points are shown from replicate experiments performed twice.

### Protein structure modeling

To investigate structural features that could explain the increased thermal stability of MDH-1 compared to MDH-2, we did protein structure homology modeling because the crystal structures of *C. elegans* MDH-1 and MDH-2 have not been determined. The MDH-1 and MDH-2 sequences were submitted to SWISS-MODEL (39), and the predicted structures were built with the crystal structures 1BMD.1.A (25) and 2DFD.1.A (unpublished structure from the Structural Genomics Consortium) as the templates, respectively. The models had GMQE and QMEAN scores (41) of 0.79 and -0.62, respectively, for MDH-1 and 8 and -0.04, respectively, for MDH-2. The MDH-1 model was refined by minimizing the free energy using the YASARA energy minimization server (42). The z-scores for the modeled MDH-1 and MDH-2 structures were -9.32 and -10.69, respectively. The bond and rotamer angles were checked using MolProbity (45, 46). The MDH-1 model had 95% favored rotamers, 98% favored Ramachandran angles, and no Ramachandran outliers. The MDH-2 model had 97% favored rotamers, 97% favored Ramachandran angles, and no outliers.

Both models passed the Verify 3D assessment (47–49). All of these results indicated that both model structures were of high quality. (The raw output from these tests are shown in supplementary Figures S3-S5, and the PDB files of the MDH-1 and MDH-2 protein structure models are posted with the supplementary information.)

Previous studies have found that more stable MDH enzymes contained more intersubunit hydrogen bonds and ion pairs in comparison to less stable ones. In addition, salt bridges (hydrogen-bonded ion pairs) may contribute more to protein stability at high temperatures (25, 65). Therefore, we searched for predicted hydrogen bonds between the subunits of both modeled MDH-1 and MDH-2 structures and focused on the salt bridges (Tables 4 and 6). We found that MDH-1 had more predicted ionic interactions between the two subunits than *S. scrofa* MDH1 but fewer interactions than the thermostabile *T. thermophilus* MDH. MDH-1 had predicted salt bridges between Asp25 and Arg22, Asp25 and Lys248 (Figure 7A), and Glu50 and Lys 236 (Figure 7D). *S. scrofa* MDH1 doesn’t form these salt bridges because it has different amino acids at these locations.

**Figure 7.**
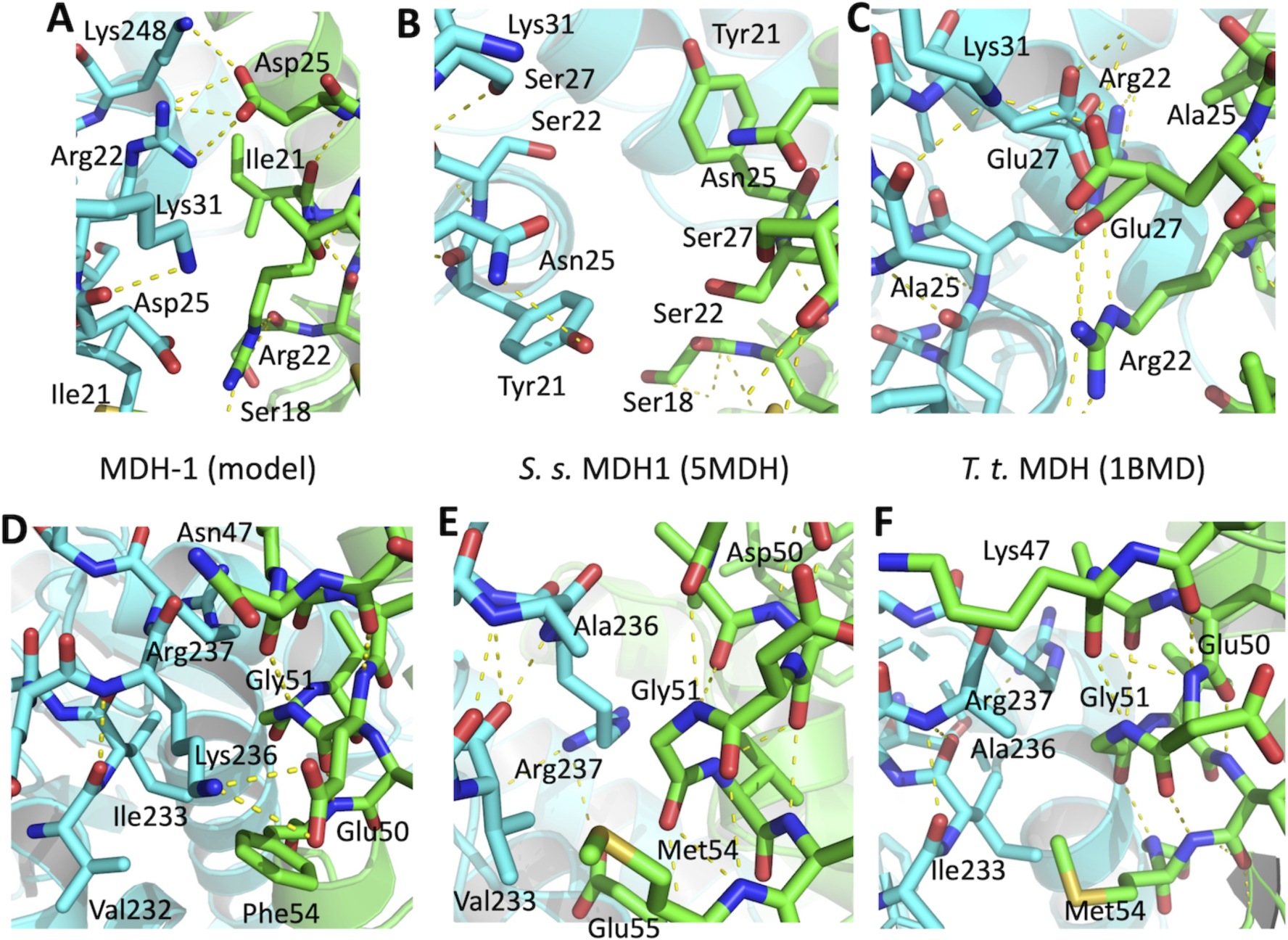
Intersubunit ionic interactions predicted in MDH-1. PyMOL was used to visualize the predicted novel ionic hydrogen bonds in the MDH-1 structural model (based on the 1BMD structure), and the corresponding views of *S. scrofa* MDH1 (5MDH) and *T. thermophilus* MDH (1BMD). Hydrogen bonds are indicated by the dashed yellow lines. Subunit A is shown in green and subunit B in teal. Amino acid numbering corresponds to the *S. scrofa* MDH1 sequence shown in Figure 1A. **A.** The hydrogen bonds between Asp25 and Arg22 and Lys248 are shown in MDH-1 with the surrounding amino acids. **B.** MDH1 has Asn25 and Ser22, which are not close enough to form a hydrogen bond. **C.** *T. thermophilus* MDH has a different, novel ionic interaction in this location between Glu27 and Lys31. (The 1BMD structure file shows Glu27 in two equally likely conformations (25)). This enzyme has an Ala25. **D.** The hydrogen bonds between Glu50 and Lys236 in MDH-1 are shown. **E.** MDH1 has an Ala236, so the same hydrogen bond can’t form. Instead, this view shows a different hydrogen bond, Glu55 to Arg237, that is shared with MDH-1 but not visible in D. **F.** *T. thermophilus* MDH also has an Ala236, and it has no predicted intersubunit hydrogen bonds in this view.

**Table 4.**
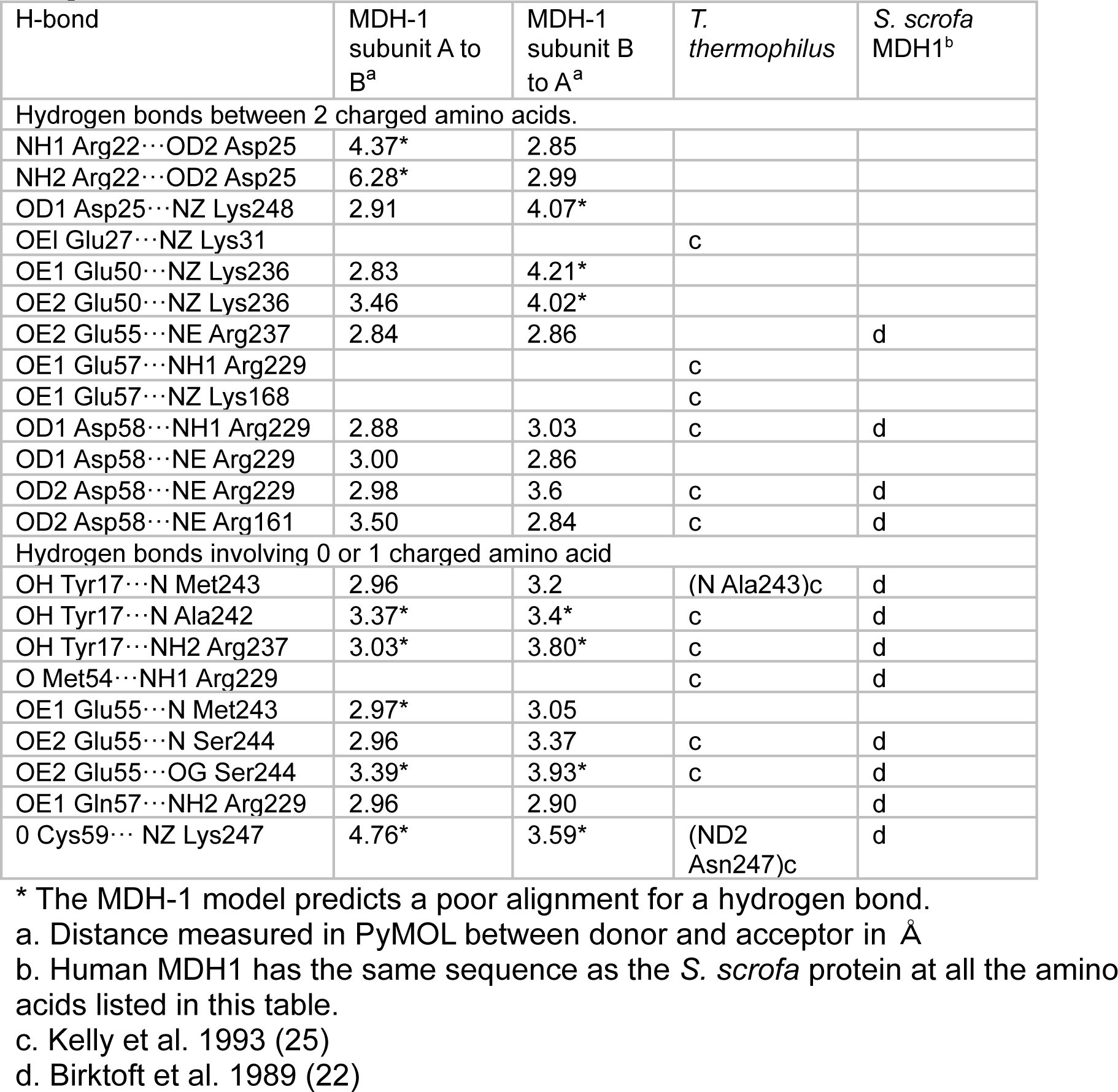
Intersubunit hydrogen bonds in MDH-1 and homologous MDH enzymes

| H-bond | MDH-1<br>subunit A to<br>B<sup>a</sup> | MDH-1<br>subunit B<br>to A<sup>a</sup> | *T.*<br>*thermophilus* | *S. scrofa*<br>MDH1<sup>b</sup> |
| --- | --- | --- | --- | --- |
| Hydrogen bonds between 2 charged amino acids. |  |  |  |  |
| NH1 Arg22···OD2 Asp25 | 4.37* | 2.85 |  |  |
| NH2 Arg22···OD2 Asp25 | 6.28* | 2.99 |  |  |
| OD1 Asp25···NZ Lys248 | 2.91 | 4.07* |  |  |
| OE1 Glu27···NZ Lys31 |  |  | c |  |
| OE1 Glu50···NZ Lys236 | 2.83 | 4.21* |  |  |
| OE2 Glu50···NZ Lys236 | 3.46 | 4.02* |  |  |
| OE2 Glu55···NE Arg237 | 2.84 | 2.86 |  | d |
| OE1 Glu57···NH1 Arg229 |  |  | c |  |
| OE1 Glu57···NZ Lys168 |  |  | c |  |
| OD1 Asp58···NH1 Arg229 | 2.88 | 3.03 | c | d |
| OD1 Asp58···NE Arg229 | 3.00 | 2.86 |  |  |
| OD2 Asp58···NE Arg229 | 2.98 | 3.6 | c | d |
| OD2 Asp58···NE Arg161 | 3.50 | 2.84 | c | d |
| Hydrogen bonds involving 0 or 1 charged amino acid |  |  |  |  |
| OH Tyr17···N Met243 | 2.96 | 3.2 | (N Ala243)c | d |
| OH Tyr17···N Ala242 | 3.37* | 3.4* | c | d |
| OH Tyr17···NH2 Arg237 | 3.03* | 3.80* | c | d |
| O Met54···NH1 Arg229 |  |  | c | d |
| OE1 Glu55···N Met243 | 2.97* | 3.05 |  |  |
| OE2 Glu55···N Ser244 | 2.96 | 3.37 | c | d |
| OE2 Glu55···OG Ser244 | 3.39* | 3.93* | c | d |
| OE1 Gln57···NH2 Arg229 | 2.96 | 2.90 |  | d |
| O Cys59··· NZ Lys247 | 4.76* | 3.59* | (ND2<br>Asn247)c | d |

\* The MDH-1 model predicts a poor alignment for a hydrogen bond.
a. Distance measured in PyMOL between donor and acceptor in Å
b. Human MDH1 has the same sequence as the *S. scrofa* protein at all the amino acids listed in this table.
c. Kelly et al. 1993 (25)
d. Birktoft et al. 1989 (22)

**Table 5.** Amino acid percentage in MDH-1 and homologous MDH enzymes

| Amino Acid | MDH-1 | *S. s.* MDH1 | *T. t.* MDH |
| --- | --- | --- | --- |
| Ala | 13.1* | 9.3 | 14.7 |
| Gly | 8.9 | 6.6 | 8.3 |
| Pro | 3.9 | 3.9 | 4.9 |
| Phe | 5.1 | 3.3 | 3.1 |
| Trp | 0.9 | 1.2 | 1.2 |
| Tyr | 1.8 | 2.1 | 2.1 |
| Ser + Thr | 11.3 | 11.7 | 7.0 |

\*The percentage of the amino acid(s) listed compared to the total number of amino acids is shown for each protein.

**Table 6.** Inter-subunit hydrogen bonds in MDH-2 and homologous MDH enzymes^a^

| Hydrogen bond | MDH-2 |  | *S. scrofa* MDH2 |  | human MDH2 |  |
| --- | --- | --- | --- | --- | --- | --- |
|  | A to B | B to A | A to B | B to A | A to B | B to A |
| OE2 Glu48...NH1 Arg233 | 3.52<br>(Asp<sup>b</sup>) | 3.65*<br>(Asp<sup>b</sup>) | 2.72<sup>c</sup> | 2.87<sup>c</sup> | 2.89 | 2.92 |
| NH1 Arg50...OD1 Asp164 | - | - | 2.90 | 2.75 | 6.32*<br>(Lys<sup>d</sup>) | 4.80*<br>(Lys<sup>d</sup>) |
| OD2 Asp43...NZ Lys217 | 2.70 | 2.58 | 2.82<sup>c</sup> | 2.94<sup>c</sup> | 3.01 | 3.44 |

\* Model predicts a poor alignment for a hydrogen bond.
a. Distance between donor and acceptor atoms in the two subunits in Å
b. MDH-2 has an Asp instead of a Glu at position 48.
c. Gleason et al. 1993 (23)
d. Human MDH2 has a Lys instead of an Arg at position 50.

Specifically, it has Asn25 & Ser22 (Figure 7B) and Asp50 & Ala236 (Figure 7E) in the corresponding positions. Figure 7E also illustrates a salt bridge in *S. scrofa* MDH1 that is also found in MDH-1 between Glu55 and Arg237 (see Table 4), but this bond is not visible in Figure 7D. *T. thermophilus* MDH has a different, novel salt bridge between Arg22 and Glu27 (Figure 7C), and it has an Ala236, so there are no intersubunit hydrogen bonds in the view shown in Figure 7F. *T. thermophilus* MDH also has an interdomain salt bridge (Arg149-Glu276) that may contribute to its stability, and this interaction is not found in *S. scrofa* MDH1 (25).

MDH-1 has Lys149 and Ser276, but there is a predicted hydrogen bond between NZ Lys149 and OD2 Asp273 in subunit B with a distance of 2.91 Å between the donor and acceptor atoms. In subunit A, these two atoms are 4.15 Å apart, and Asp273 makes hydrogen bonds with Ser276 instead. The structures of the two subunits in *S. scrofa* MDH1 and MDH2 are quite similar to each other but not identical (22, 23), and this is the source of the disparity between the hydrogen bond lengths in the two subunits.

In addition to ionic interactions, other protein structural features can contribute to protein stability, and different enzymes use different strategies to stabilize their structures (66–68). For example, some thermostable MDH enzymes have a high percentage of alanine residues and aromatic residues (69). Studies of other proteins, such as T4 lysozyme and alcohol dehydrogenase, have found that more thermostable proteins have more prolines and fewer glycines (70, 71). The *T. thermophilus* MDH sequence follows some of these trends (Table 5). *T*.

*thermophilus* MDH has a large percentage of Ala and Pro residues, an intermediate percentage of Gly residues, and the same percentage of aromatic residues as *S. scrofa* MDH1. MDH-1 has more Ala and Phe residues but also more Gly residues and the same number of Pro residues compared to *S. scrofa* MDH1. Therefore, MDH-1 has some but not all of the stabilizing amino acid characteristics found in other thermostable proteins. *T. thermophilus* MDH has fewer Ser and Thr residues than the other two proteins. Another study found that thermostabile MDH enzymes had larger volumes than mesophilic enzymes (66). Using the MDH-1 model and the crystal structures of *S. scrofa* MDH1 and *T. thermophilus* MDH, we measured volumes of 80,600, 83,600, and 81,200 Å^3^, respectively. Therefore, these proteins don’t follow the predicted trend in volumes since *S. scrofa* had the largest volume.

When we searched for predicted ionic interactions in the MDH-2 model, we found fewer interactions than in *S. scrofa* MDH2 (Table 6). We also used the unpublished human MDH2 structure (4WLU) to check for hydrogen bonds.

Gleason et al. (1993) mentioned two intersubunit hydrogen bonds in *S. scrofa* MDH2, between Glu48 and Arg233 and between Asp43 and Lys217 (23). The interaction between Asp43 and Lys217 appeared to be conserved in all three proteins. The Glu48-Arg233 interaction was conserved in human MDH2, but the bond length in MDH-1 was longer because MDH-2 has an Asp48. Our analysis of *S scrofa* MDH2 predicted another hydrogen bond between Arg50 and Asp164, which was not conserved in MDH-2, and it probably doesn’t form in human MDH2 because the human protein has a Lys50, and its terminal amino group is quite distant from Asp164.

## Discussion

We performed enzyme kinetics, thermostability, and protein structure analyses of *C. elegans* MDH-1 and MDH-2. These enzymes have not been characterized before, even though they are central metabolic enzymes in an important experimental organism. *C. elegans* is not a thermophilic organism, but it can withstand a variety of environmental conditions. These conditions could cause selection pressure on the *C. elegans* metabolic enzymes.

### MDH-1 and MDH-2 thermal stability

A larger number of salt bridges between subunits has been shown previously to increase the thermal stability of enzymes such as glutamate dehydrogenase and MDH (25, 69, 72, 73). We found that homology modeling of *C. elegans* MDH-1 predicted more intersubunit salt bridges and ion pairs compared to *S. scrofa* MDH1 but fewer compared to *T. thermophilus* MDH (Table 4). If these interactions contributed to the thermostability of MDH-1, we would expect MDH-1 to be more thermostabile than *S. scrofa* MDH1 but less thermostable than *T. thermophilus* MDH. We found that *C. elegans* MDH-1 lost 50% of its activity when incubated at 55 °C for about 17 min in 0.5 mM NADH and 50 mM Tris, pH 7.5 (Figure 6). Trejo et al. (2001) measured a half time of inactivation for His-tagged *S. scrofa* MDH1 of 13 min at 55 °C in 50 mM sodium phosphate buffer, pH 7.4 (73). A previous study measured a half time of inactivation of endogenous *S. scrofa* MDH1 of 20 min at 55 °C in 0.1 M phosphate buffer, pH 7.4 (74). *T. thermophilus* MDH remained fully active after incubation at 90 °C for 60 min in 20-33 mM potassium phosphate, pH 7.0 (75, 76). Another study measured a half time of inactivation of *T. thermophilus* MDH of 34 min at 90 °C in 20 mM phosphate buffer, pH 7.4 (77). Therefore, it appears that MDH-1 had a similar thermostability as *S. scrofa* MDH1, and MDH-1 was significantly less thermostable than *T. thermophilus* MDH. Perhaps the additional intersubunit ionic interactions in *C. elegans* MDH-1 optimize enzyme activity in other conditions besides high temperatures.

We observed the highest enzyme activity for MDH-1 and MDH-2 at 40 and 35 °C, respectively (Figure 5). Recombinant His-tagged *S. scrofa* MDH1 had maximum activity at 60 °C (73). This difference seems reasonable since *C. elegans* have an optimum growing temperature around 20 °C, while mammals typically maintain a 37 °C temperature. Even though MDH-1 had lower activity at reaction temperatures above 40 °C, the enzyme could retain activity for about 10 min. when it was heated to 55-57°C and then quickly cooled and assayed at 24°C (Figure 6). This suggests that the denaturation of MDH-1 may be reversible up to a point if the enzyme lost activity in assays performed above 40 °C because it started to denature. The ability to refold is a feature of some thermostable enzymes and may be advantageous for maintaining activity under adverse conditions (66). If an enzyme refolds incorrectly, a buried, charged residue mis-aligned from its binding partner could be very unstable. These higher energy folding intermediates may facilitate native enzyme refolding by preventing enzymes with numerous buried salt bridges, like MDH-1, from being trapped in a low-energy, non-native conformation (65).

We measured a rate of inactivation of MDH-2 of 2.85 min^-1^ (which corresponds to a t_1/2_ of 0.24 min) under the same conditions as those used for MDH-1 (Figure 6), and homology modeling predicted fewer intersubunit ion pairs compared to *S. scrofa* MDH2 (Table 6). One study reported a t_1/2_ of inactivation of *S. scrofa* MDH2 at 60 °C in 20 mM phosphate buffer, pH 7.4, of 1.8 min (77), suggesting that *S. scrofa* MDH2 was more stable than MDH-2 since MDH-2 lost activity faster at a lower temperature. These results are consistent with the difference in intersubunit interactions in the two enzymes. In addition, based on our analysis of the human MDH2 structure (Table 6), we would predict that it is intermediate in stability between MDH-2 and *S. scrofa* MDH2.

MDH-2 may be structurally less stable because MDH2 enzymes have been found to bind to fumarase and citrate synthase in the mitochondria (reviewed in (78, 79)). This facilitates substrate channeling through the citric acid cycle because the forward reaction of malate dehydrogenase is thermodynamically unfavorable while the citrate synthase reaction is favorable (61). The binding between *S. scrofa* MDH2 and citrate synthase was found to be specific but transient (80). When we purified endogenous MDH-2, it is likely that MDH-2 was separated from fumarase and citrate synthase. The interactions between MDH-2 and its binding partners may stabilize the structure of this protein complex so that it can function even if the separate enzymes are not as stable.

The stability differences between MDH-1 and MDH-2 may partly be the result of their different phylogenetic origins. Numerous studies have shown that MDH enzymes cluster into three clades (1, 81, 82), and two of these clades are evident in Figure 1. The sequences of mitochondrial MDH enzymes have a high sequence similarity to some eubacterial MDH enzymes, consistent with the theory of a bacterial origin of mitochondria (83, 84). The ancestors of eukaryotic cells are still an active area of research, but recently archaeal genomes have been sequenced that contain classes of proteins considered to be eukaryotic signature proteins, such as the phylum Lokiarchaeota (85, 86). However, this second clade of archaeal MDH enzymes are more similar to lactate dehydrogenases than to MDH enzymes (1, 81, 82). The third clade of MDH enzymes include other eubacterial enzymes, such as *T. thermophilus*, and cytoplasmic MDH1 enzymes, and these are highly diverged from the mitochondrial MDH2 enzymes (2, 76) Perhaps the bacterial ancestors of cytoplasmic MDH were more stable. Interestingly, additional thermostable bacterial MDH enzymes have been found that were more similar to cytoplasmic MDH than mitochondrial MDH, such as those from *Streptomyces coelicolor* A3(2) and *Streptomyces avermitilis* MA-4680 (87, 88).

### MDH-1 and MDH-2 functions in *C. elegans* dauer larvae

In our analysis of the intersubunit ionic interactions for MDH-1 and MDH-2, we found that MDH-1 had more intersubunit interactions compared to the homologous mammalian enzymes, while MDH-2 had fewer. This suggested that there were differences in the selective pressures against each enzyme. We hypothesized that one reason for the increased stability of MDH-1 was because it was important for metabolism in dauer larvae, while the requirement for MDH-2 was reduced. There is support for this in the literature. Dauer larvae can be produced naturally, such as by increasing the *C. elegans* density and reducing food availability, or artificially using mutant strains. The *daf-2* strains have mutations in an insulin receptor, and different strains form dauer larvae either constitutively or in a temperature-dependent manner (15, 89). Strains with *daf-2* mutations have an extended lifespan (90), and they have a similar, but not identical, metabolic signature as dauer worms (16, 91). Whole genome microarray experiments showed that the *mdh-1* gene was overexpressed in dauer larvae while *mdh-2* was underexpressed (16, 21, 92). Another study used serial analysis of gene expression and found that *mdh-2* was underexpressed in dauer larvae (93). Fuchs et al. (2010) reported an increase in expression of *mdh-1* in *daf-2* dauer worms (91). The protein MDH-1 was more abundant in *daf-2* worms compared to control worms (94, 95) and in *daf-2* dauers grown at 25 °C compared to *daf-2* larvae grown at the permissive temperature (96). However, Depuydt et al. (2014) found that levels of MDH-2 protein were slightly elevated in *daf-2* worms (94). The fact that MDH-1 expression was elevated in dauer worms suggests that it plays an important role in this life cycle stage.

MDH-1 expression is likely elevated in dauer larvae because the metabolic processes occurring in dauers depend on its activity. There are three main pathways that are relevant, and they have each been emphasized in different studies. The first pathway involves the glyoxylate shunt, which is important in dauer larvae since they rely on stored lipids for energy. As discussed above, malate produced in the mitochondria by the glyoxylate shunt bypasses MDH-2 and is exported to the cytoplasm and converted to oxaloacetate by MDH-1. The expression of genes involved in the glyoxylate shunt are up-regulated in dauer larvae (16, 21). In *daf-2* dauer worms, the main glyoxylase enzyme, ICL-1, is up-regulated when measured by mass spectrometry (96). A study of enzyme activities in protein extracts derived from adult worms or dauer larvae found that the activity of isocitrate dehydrogenase was greatly reduced in dauer larvae compared to adult worms, but the activity of isocitrate lyase was only slightly reduced in dauers, suggesting that a larger fraction of isocitrate was metabolized via the glyoxylate pathway compared to the citric acid cycle in dauer worms (19).

Therefore, the importance of the glyoxylate shunt in dauer metabolism is well supported.

The second relevant pathway in dauers is gluconeogenesis. Oxaloacetate produced by MDH-1 in the cytoplasm is converted to phosphoenolpyruvate (PEP) by PEP carboxykinase (PEPCK), and PEP can be used for gluconeogenesis. There has been some debate in the literature about how much gluconeogenesis actually occurs in dauer larvae. One would expect gluconeogenesis to occur since dauer larvae do not feed. Also, numerous studies have shown that proteins involved in gluconeogenesis, such as PEPCK, are up-regulated in dauer larvae, and this has been shown using gene expression or protein levels measured by mass spectrometry (16, 21, 96).

However, in a study using enzyme assays in protein extracts, the authors found that PEPCK activity was similar in dauer larvae compared to adult worms, and fructose 1,6-bisphosphatase activity was low in dauers (97). Since fructose 1,6-bisphosphatase is an important regulatory step in gluconeogenesis, the authors argued that it was more likely that PEPCK was functioning in reverse to fix CO_2_ and form oxaloacetate. In other words, high PEPCK levels did not indicate the predominant direction of the PEPCK reaction. Under specific environmental conditions, such as mild desiccation (98% relative humidity), *daf-2* dauer larvae are able to initiate gluconeogenesis to make trehalose, and the amount of trehalose formed depended on the presence of the *icl-1* gene, which encodes one of the main glyoxylase shunt enzymes. However, when not exposed to mild desiccation, dauer larvae contained lower levels of trehalose that were similar whether *icl-1* was present or not (14). Therefore, dauer larvae were capable of up-regulating gluconeogenesis when necessary, and this required the glyoxylate pathway and would require MDH-1, but dauers did not seem to carry out gluconeogenesis constitutively.

A third metabolic process in dauer worms called malate dismutation can also involve MDH-1. Malate can be generated by PEPCK and MDH-1 in the cytoplasm and transported into the mitochondria (94, 97), or malate can be produced by the glyoxylate shunt in the mitochondria. There, malate can be metabolized via malate dismutation, which requires specialized enzymes (see Figure S6 for a diagram). In this process, some of the malate is converted to fumarate by fumarase, while some of the malate is converted to pyruvate via malic enzyme, a process that generates NADH. The NADH provides reducing equivalents to Complex I to reduce rhodoquinone, an electron carrier similar to ubiquinone, which is used by fumarate reductase (an alternative version of Complex II) to convert fumarate to succinate (98, 99). A few electrons are transported across the inner mitochondrial membrane by Complex I to generate 21 ATP via oxidative phosphorylation, but the final electron acceptor is fumarate rather than oxygen, so dauer worms consume less oxygen (14, 100).

Interestingly, malate dismutation appears to occur in many long-lived mutant strains of *C. elegans*, and oxaloacetate and malate can extend the lifespan of *C. elegans* (94, 101–104). However, the specific processes required for life-span extension are varied, and they are actively discussed in the literature. During malate dismutation, MDH-1 acts in the energetically favorable direction (from oxaloacetate to malate), while MDH-2 is not involved in this pathway.

Other processes occur in dauer larvae to maintain enzyme activity at higher temperatures. For example, in wild-type and *daf-2* worms, the presence of exogenous or endogenously produced trehalose increased the thermotolerance of the worms (105, 106). Trehalose has also been found to increase the thermal stability of enzymes in vitro (107). In *Saccharomyces cerevisiae*, trehalose stabilized proteins during heat shock and reduced aggregation of partly unfolded proteins, but trehalose was rapidly degraded after heat shock to allow proteins to refold (108). Similarly, trehalose could stabilize proteins in *C. elegans* and reduce protein aggregation (105, 109). The presence of trehalose could cause greater thermostability of enzymes in dauer larvae and assist with protein refolding.

Therefore, multiple mechanisms may allow proteins to function under adverse conditions in dauer larvae.

### MDH-1 and MDH-2 functions during favorable growth conditions

It seems less likely that the differential stability of MDH-1 and MDH-2 result from reactions occurring during favorable, aerobic growth conditions because both MDH-1 and MDH-2 are required. Specifically, both enzymes act in the malate-aspartate shuttle to transport electrons across the inner mitochondrial membrane. Since the inner membrane is impermeable to NAD^+^ and NADH, other molecules are used to transport reducing equivalents to the electron transport chain. In this process, NADH from glycolysis is used by MDH-1 to convert oxaloacetate to malate, which is transported across the inner membrane via the 2-oxoglutarate/malate carrier protein, called MISC-1 in *C. elegans* (110). Inside the mitochondrial matrix, MDH-2 regenerates the NADH. The resulting oxaloacetate can either enter the citric acid cycle or be transaminated to aspartate and then transported back out of the matrix via a calcium-binding mitochondrial carrier protein, which is human SLC25A12 and probably K02F3.2 in *C. elegans* (111, 112). This process is sustainable because 2-oxoglutarate is exported out of the matrix by MISC-1, and aspartate is used to transaminate 2-oxoglutarate to glutamic acid in the inner membrane space. This regenerates oxaloacetate in the cytoplasm. Also, glutamic acid can return to the matrix via K02F3.2 and be the amino group donor for transaminases. This process requires the activity of both MDH enzymes and both antiporter carrier proteins.

### Enzyme Kinetics Results

Our enzyme kinetics results for MDH-1 and MDH-2 were similar to those found for homologous enzymes. Previous studies have reported *K*_M_ values for *S. scrofa* MDH1 and MDH2 of 30-35 μM and 40 μM, respectively, under similar reaction conditions and a k_cat_ value of 475 s^-1^ for *S. scrofa* MDH1 (73, 113). Our results were also similar to those found for recombinant cytoplasmic MDH from the tapeworm *Taenia solium*. This enzyme had a *K*_M_ for oxaloacetate and NADH of 44 μM and 167 μM, respectively, and a specific activity of 720 Units/mg protein (6). More divergent results were found for recombinant cytosolic MDH from the liver fluke *Fasciola gigantica*. This enzyme had a *K*_M_ for oxaloacetate and NADH of 276 μM and 141 μM, respectively (5). Other MDH enzymes from invertebrates had a wide range of *K*_M_ and specific activity values, but for most of these studies, the results couldn’t be directly compared to our results because the reaction conditions were too different.

The enzyme kinetics constants for MDH-1 and MDH-2 were similar to each other except for the *K*_M_ of NADH (Figure 4C). Some researchers have compared the kinetics of *S. scrofa* MDH1 and MDH2. They did not determine *K*_M_ values, so their results are not directly comparable to the experiments shown in Figure 4.

However, they did do simulations showing that MDH1 had a higher enzymatic activity than MDH2 at low [NADH] (62), and this is consistent with our measurement of a lower *K*_M_ for NADH for MDH-1 compared to MDH-2 (Figure 4C). The lower *K*_M_ for MDH-1 would also allow it to have more activity at the lower [NADH] in the cytoplasm compared to the mitochondrial matrix (reviewed in (114)).

## Data availability

All data are contained within the manuscript and Supporting Information.

## Supporting information

MDH-1 model

MDH-2 model

## Acknowledgements

We are grateful to Wei Gu, Megan Gautier, Anamica Bedi, Penelope Lindsay, Alexander Strzalkowski, Sanchita Mukherjee, Robert Hincapie, Valeria Valbuena, and Ana Cheng. They contributed to various parts of this project, especially constructing the protein expression plasmids and growing *C. elegans*.

## Authors’ Contributions

MJT, ERC, DSR, and KMW designed and performed the experiments. ERC, DSR, and KMW performed computational analyses. KMW wrote the manuscript with assistance from MJT, ERC, and DSR. KMW supervised the project and assembled the figures.

## Footnotes

1 Abbreviations: MDH, malate dehydrogenase in general or bacterial MDH; MDH-1, *C. elegans* cytoplasmic MDH-1; MDH-2, *C. elegans* mitochondrial MDH-2; MDH1, mammalian cytoplasmic MDH1; MDH2, mammalian mitochondrial MDH2; *mdh-1*, the gene encoding MDH-1; *mdh-2*, the gene encoding MDH-2

