## Supporting Information for "Kinetic characterization and thermostability of *C. elegans* cytoplasmic and mitochondrial malate dehydrogenases"

### Contents of Supporting Information

**Supporting Experimental Procedures**

**Figure S1. Recombinant MDH-1 purification**

**Figure S2. pH-dependence of MDH-2 activity**

**Figure S3. ProSA-web z-score graphs for MDH-1 and MDH-2 homology models**

**Figure S4. Ramachandran diagrams for the MDH-1 and MDH-2 models**

**Figure S5. Verify 3D results for the MDH-1 and MDH-2 models**

**Figure S6. Diagram of malate dismutation reactions**

### Supporting Experimental Procedures

#### Alternate MDH-1 protein structure models

The MDH-1 structure was also modeled using 5MDH (116) and 4MDH (117) as templates (data not shown). These initial models had more bond and rotamer angle outliers both before and after minimization compared to the model based on 1BMD. Energy minimization results using UCSF Chimera and the YASARA server were also compared, and the YASARA server consistently produced fewer bond and rotamer angle outliers.

#### Purity of endogenous MDH-1

The *C. elegans* worms used for the endogenous protein purifications were grown on *E. coli* as the food source. The worms were washed extensively to remove the bacteria from their gut before the worms were frozen, but it is possible that some *E. coli* MDH contaminated our endogenous enzyme samples. The *E. coli* MDH enzyme is 32.3 kDa with a theoretical pI of 5.6. This pI is closer to the pI of MDH-1 than MDH-2. If the *E. coli* enzyme were present as a contaminant, it may have co-purified with MDH-1. There is a contaminating band in some of the DEAE fractions at this molecular weight (lane 8 in Figure 3A), but it is quite faint, especially in the combined DEAE fractions (lane 9 in Figure 3B). The $K_M$ value for *E. coli* MDH was found to be 40-49 µM for oxaloacetate with a specific activity of 1920 U/mg (118-120). The $K_M$ values for oxaloacetate were very similar for endogenous and recombinant MDH-1 (Figure 4), and the specific activities of the endogenous MDH-1 and MDH-2 enzymes were similar to each other, as were the specific activities of the recombinant enzymes (Tables 1 and 3). If there was significant contamination of *E. coli* MDH in our MDH-1 sample, we would expect it to lower the $K_M$ or increase the specific activity of endogenous MDH-1, and this was not observed.

### Figure S1. Recombinant MDH-1 purification

**A**

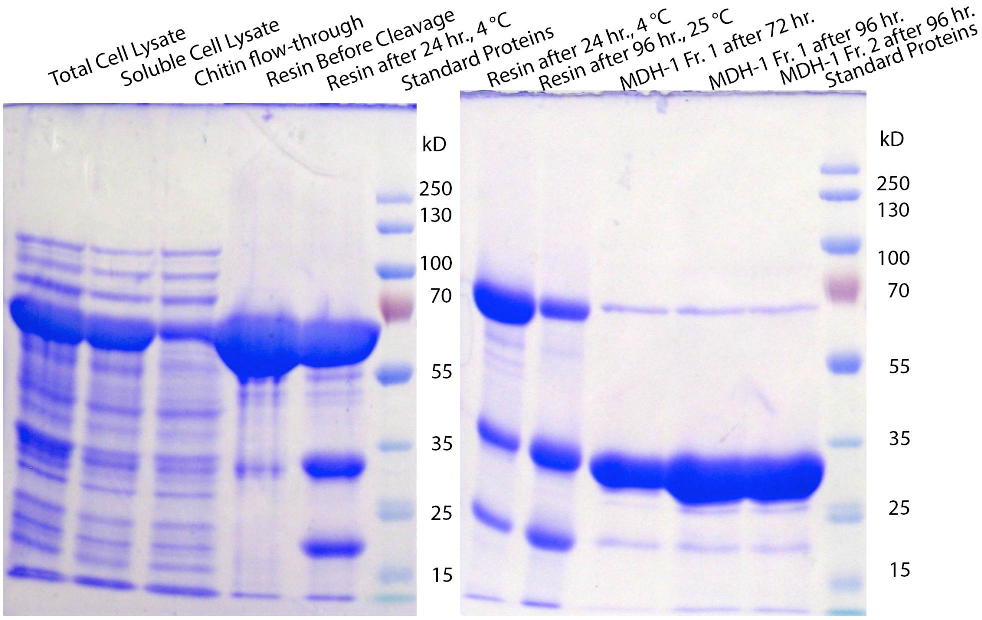


**B**

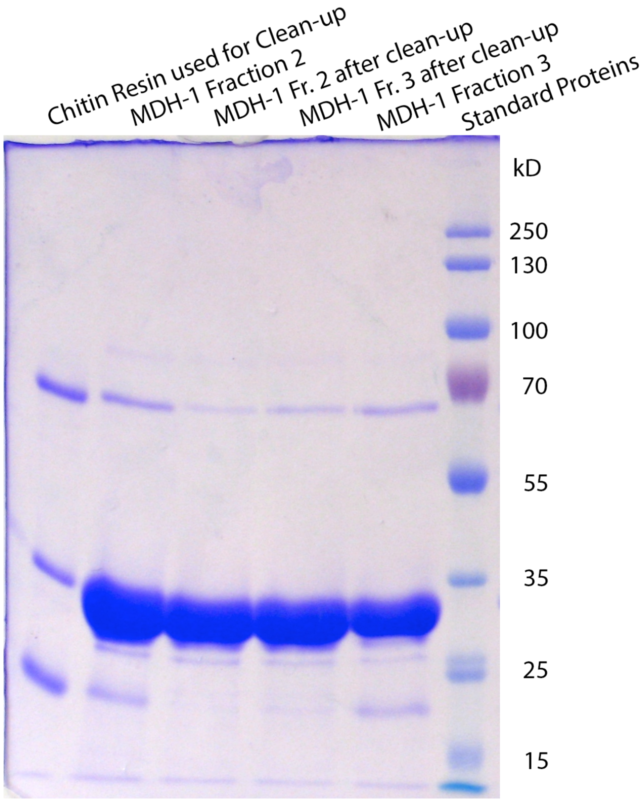

**Figure S1. Recombinant MDH-1 purification**

**A.** Samples from the purification of recombinant MDH-1 are shown in 10% SDS-PAGE gels. The soluble cell lysate containing the MDH-1-CBD fusion protein (63.7 kDa) was run through a chitin resin column, and the protein bound to the resin (lane 4). Then DTT was added to the resin for 24 hr. at 4 °C to induce cleavage of the intein to release MDH-1 (36 kDa). Since the cleavage was incomplete (lanes 5 and 7), fresh DTT was added, and the column was left at 25 °C for 96 hr (lane 8). The MDH-1 fractions (lanes 10-11) showed that the intense MDH-1 band was contaminated with MDH-1-CBD fusion protein and by cleaved CBD (20 kDa). The standard protein sizes are shown in kDa.

**B.** Examples of recombinant MDH-1 (36 kDa) samples are shown that were further purified by passing the eluted MDH-1 fractions (e.g., lane 11 in A) through a fresh chitin column to remove contaminating MDH-1-CBD fusion protein (63.7 kDa) and cleaved CBD (20 kDa). Lane 1 shows the chitin resin after it was used to clean the fractions. Lanes 2 and 5 show the original fractions 2 and 3, respectively, and lanes 3 and 4 show the same samples after they were passed through the second chitin column. The sample shown in lane 3 was used for enzyme kinetics experiments. The standard protein sizes are shown in kDa.

**Figure S2. pH-dependence of MDH-2 activity**

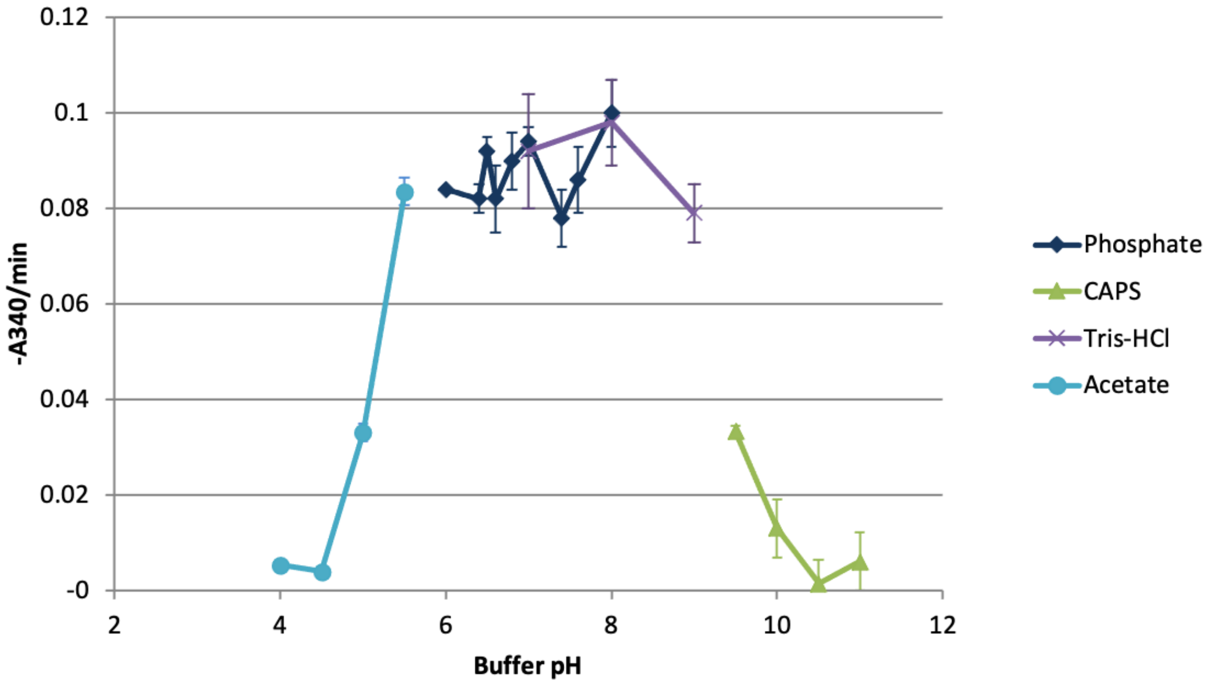


**Figure S2. pH-dependence of MDH-2 activity**
The reaction velocity in absorbance/min from a constant amount of MDH-2 was measured in buffers at the pH values shown on the X-axis. The legend shows the buffer type used in each pH range. Each buffer was at 0.1 M and contained 0.1 M KCl, 0.15 mM oxaloacetate, and 56 μM NADH. pH values were measured at 24 °C. The symbols show the mean value of 3 measurements, and the error bars show the standard deviation.

**Figure S3. ProSA-web z-score graphs for MDH-1 and MDH-2 homology models**

**A**

### Results for MDH-1_1bmd1_YASARAmin_end.pdb, chain A (331 aa)

#### Overall model quality

HELP

*Z*-Score: **-9.32**

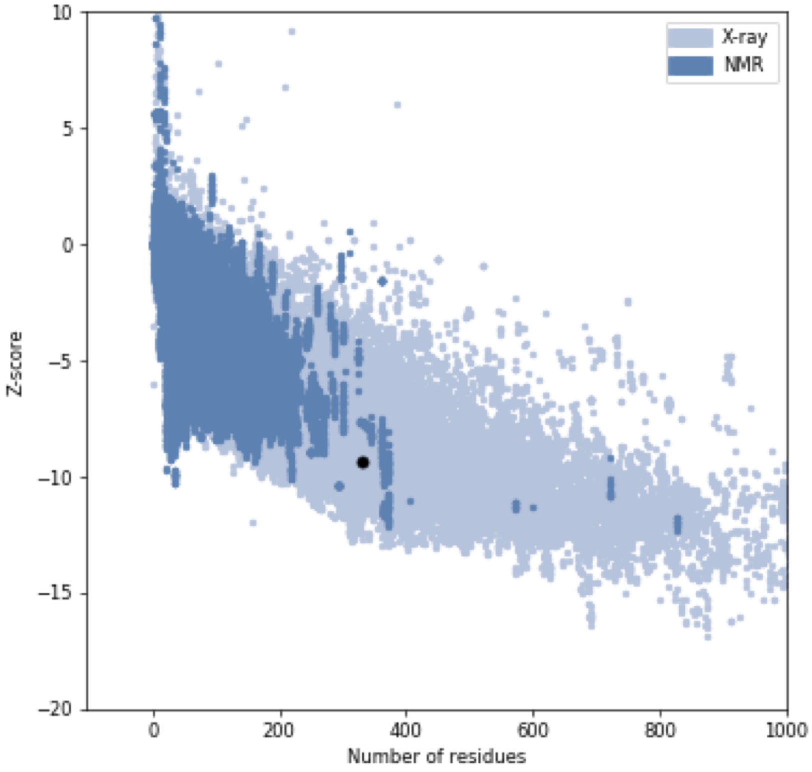

**B**

### Results for MDH-2_model_plusH_2dfd1A.pdb, chain A (313 aa)

#### Overall model quality

HELP

*Z*-Score: **-10.69**

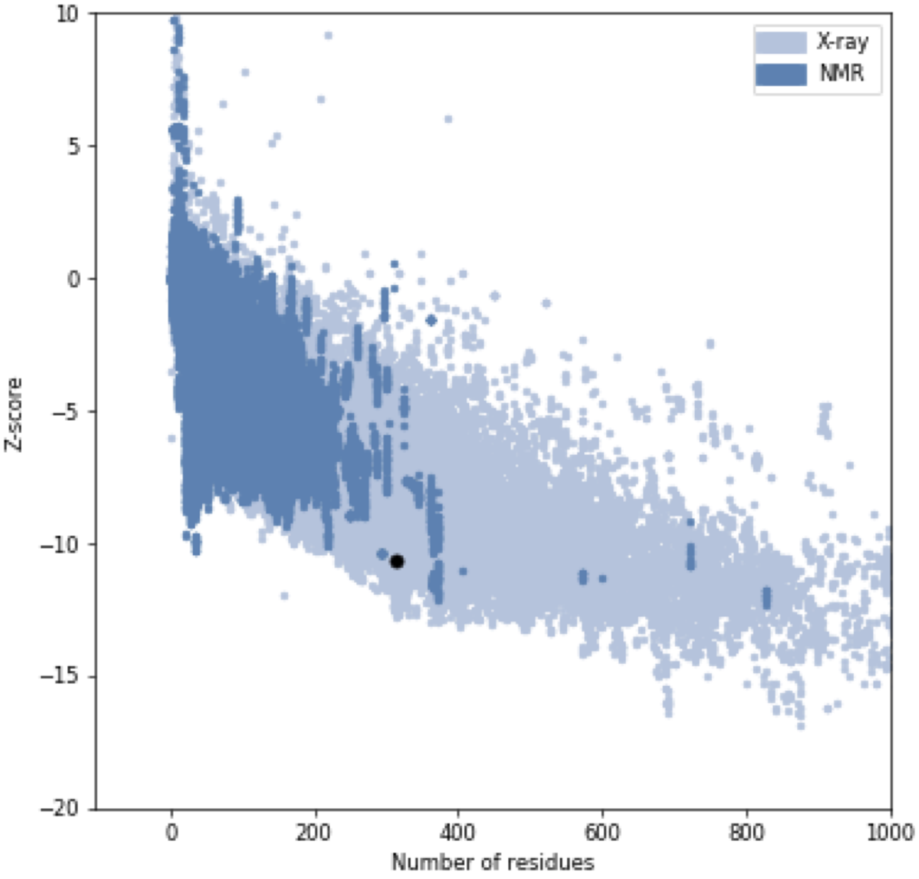


**Figure S3. ProSA-web z-score graphs for MDH-1 and MDH-2 homology models.** The homology model structures were submitted to the ProSA-web server, and the z-scores were calculated. The graphs show the z-scores for other X-ray and NMR protein structures in the PDB database, and the black dots show the scores for **A.** MDH-1 and **B.** MDH-2. Both scores are in the range of those expected for a good quality model.

### Figure S4. Ramachandran diagrams for the MDH-1 and MDH-2 models

**A**

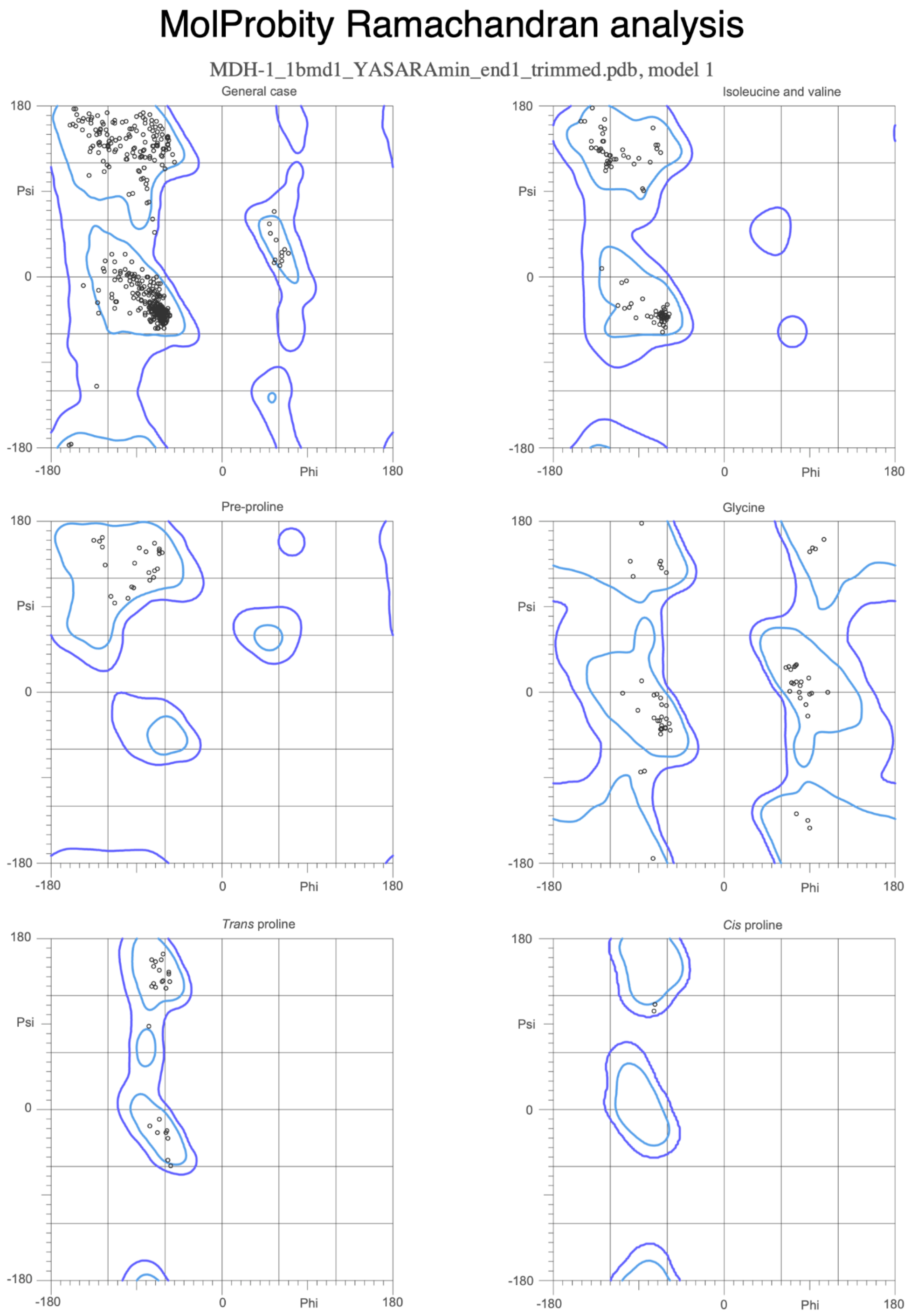


97.9% (644/658) of all residues were in favored (98%) regions.
100.0% (658/658) of all residues were in allowed (>99.8%) regions.

There were no outliers.

**B**

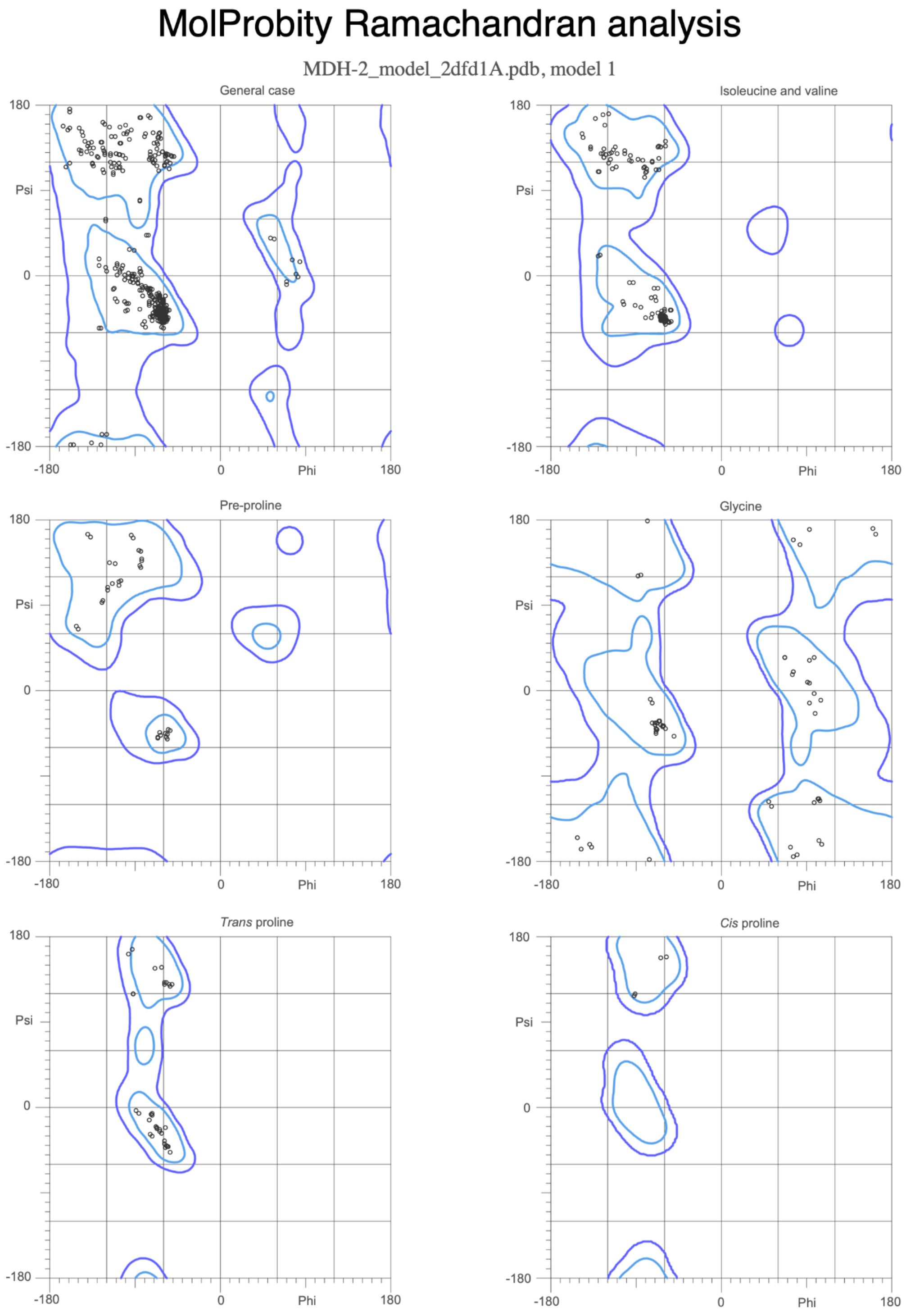


96.8% (602/622) of all residues were in favored (98%) regions.
100.0% (622/622) of all residues were in allowed (>99.8%) regions.

There were no outliers.

**Figure S4. Ramachandran diagrams for the MDH-1 and MDH-2 models**
The MDH-1 and MDH-2 structural models were submitted to the MolProbity website, and the Ramachandran diagrams were produced. **A.** The MDH-1 model had 98% favored Ramachandran angles and no Ramachandran outliers. **B.** The MDH-2 model had 97% favored Ramachandran angles and no outliers.

### Figure S5. Verify 3D results for the MDH-1 and MDH-2 models

### A

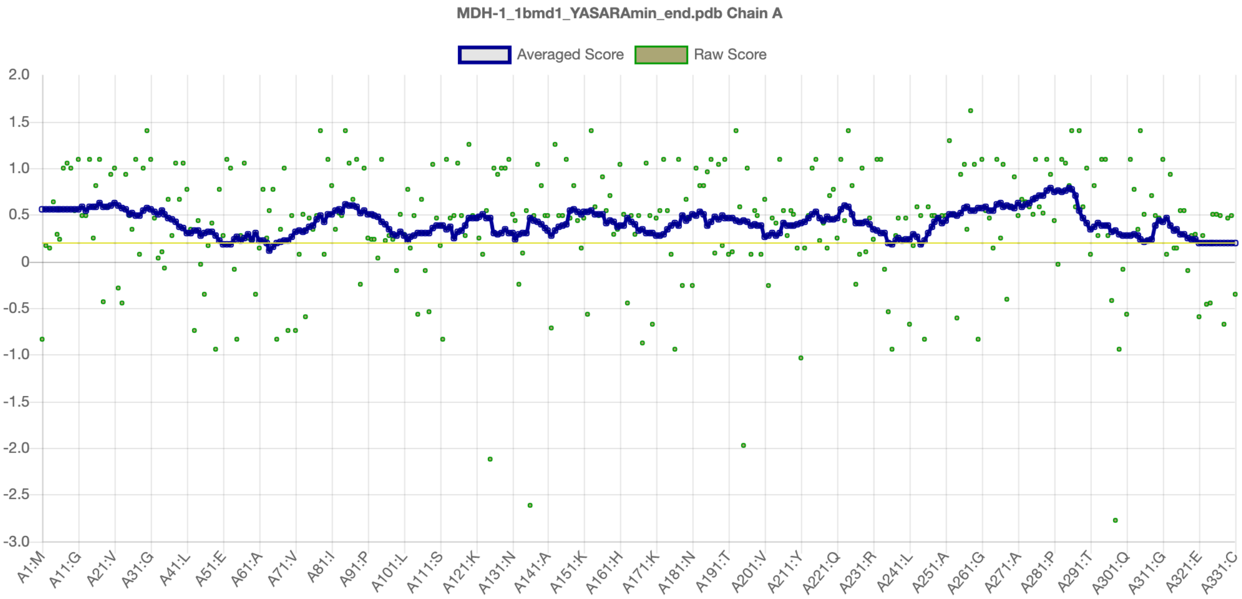


### B

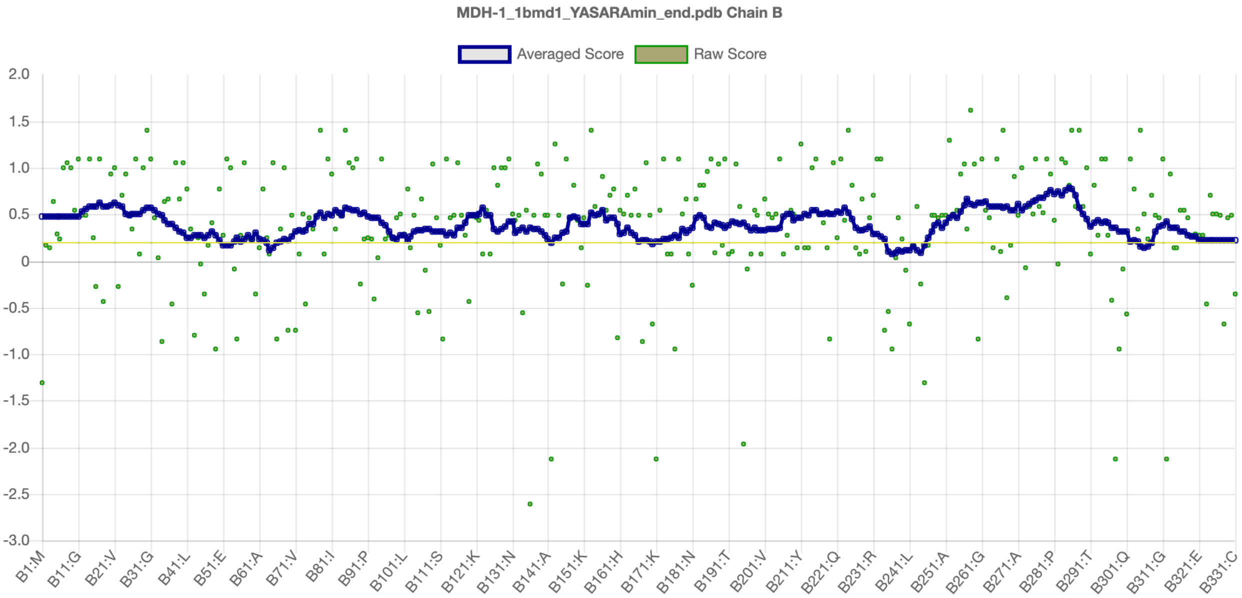

C

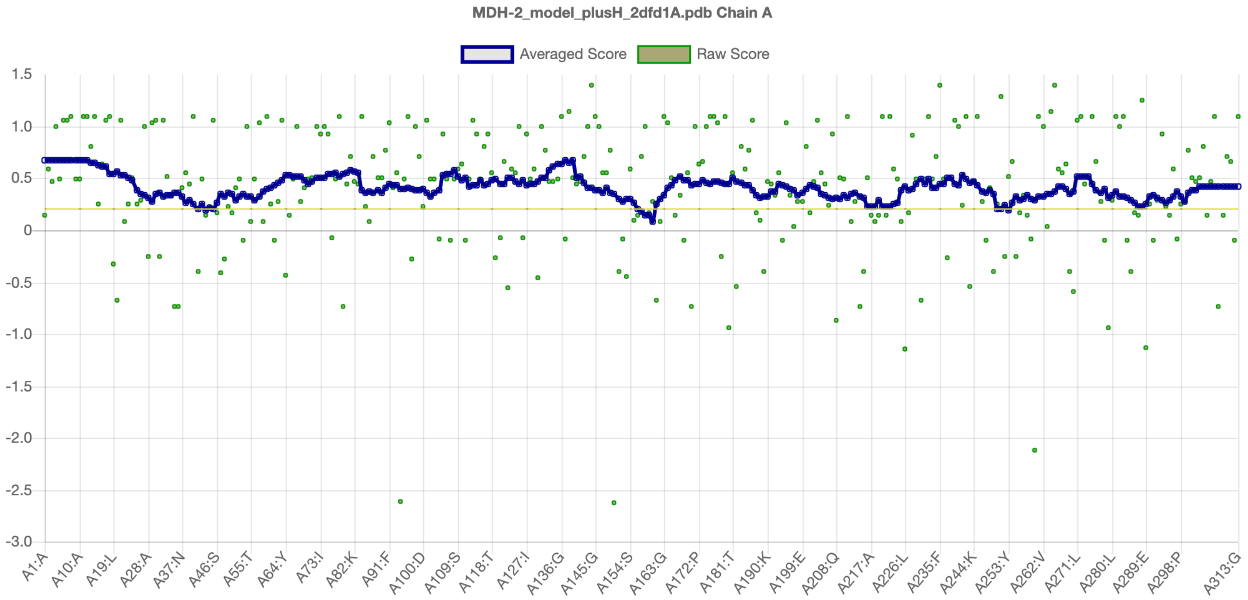


D

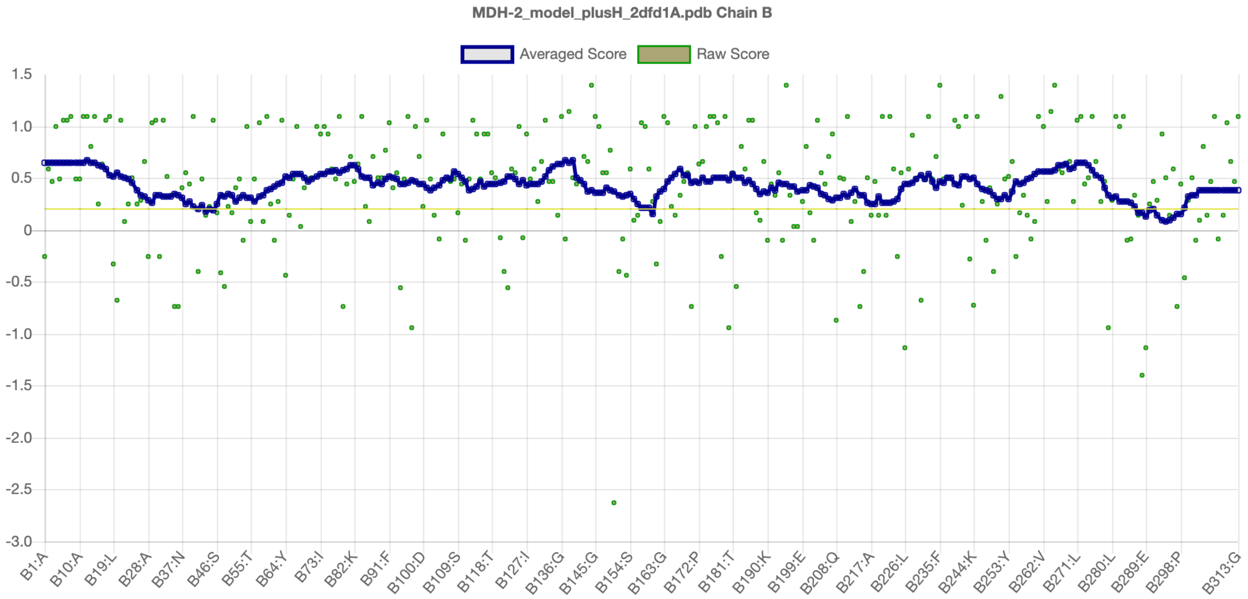


**Figure S5. Verify 3D results for the MDH-1 and MDH-2 models**

The MDH-1 and MDH-2 structural models were submitted to the SAVES v6.0 web server (https://saves.mbi.ucla.edu/), and the Verify 3D assessment was run. **A.** Plot for MDH-1, subunit A. **B.** Plot for MDH-1, subunit B. **C.** Plot for MDH-2, subunit A. **D.** Plot for MDH-2, subunit B. MDH-1 had 95.62% of the residues with an averaged 3D-1D score ≥ 0.2, and MDH-2 had 96.81% of the residues with an averaged 3D-1D score ≥ 0.2. Therefore, they both passed this assessment.

**Figure S6. Diagram of malate dismutation reactions**

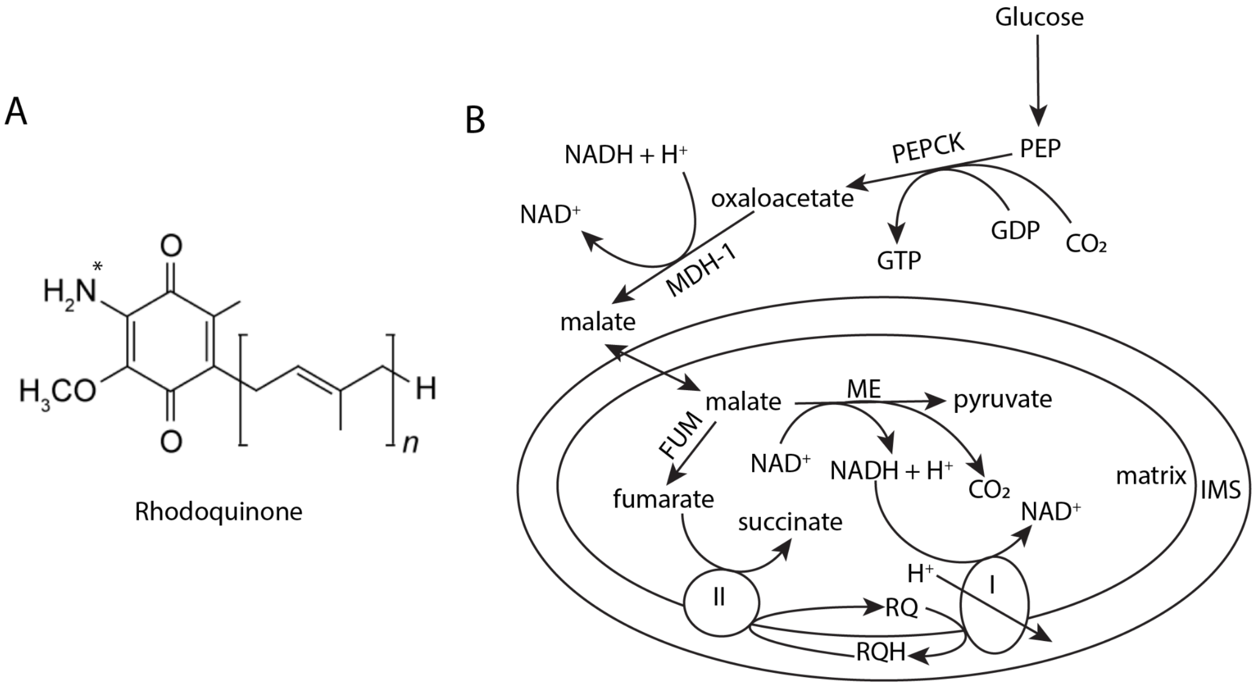


**Figure S6. Diagram of malate dismutation reactions.**
**A.** The structure of rhodoquinone. The star marks the amino group that replaces the methoxy group of ubiquinone. **B.** This diagram illustrates the reactions related to malate dismutation that are discussed in the manuscript. Glucose or glycerol can be sources of PEP. Then PEP can be converted to malate in the cytoplasm via phosphoenolpyruvate carboxykinase (PEPCK) and MDH-1. Then malate is transported into the mitochondria, where it can be metabolized by either fumarase (FUM) to form fumarate or by malic enzyme (ME) to form pyruvate. Malate can also be produced in the mitochondria via the glyoxylate shunt. The NADH produced by ME can be oxidized by Complex I, which reduces rhodoquinone (RQ) instead of ubiquinone. Then reduced RQH can be oxidized by Complex II, which is acting as a fumarate reductase here. The result is that Complex I transports electrons across the inner membrane, and succinate is the final product in this simplified and anaerobic electron transport chain. IMS is inner membrane space.
